# ENPP1 inhibitor with ultralong drug-target residence time as an innate immune checkpoint blockade cancer therapy

**DOI:** 10.1101/2025.05.18.654655

**Authors:** Randolph M. Johnson, Songnan Wang, Jacqueline A. Carozza, Daniel Fernandez, Jan Scicinski, Neil A. Verity, Rachel Mardjuki, Xujun Cao, Jacqueline Papkoff, Nigel Ray, Lingyin Li

**Author notes:** These authors contribute equally to this work (Randolph M. Johnson, Songnan Wang). Corresponding author: Lingyin Li.

## Abstract

Only one in five patients is estimated to respond to immune checkpoint inhibitors, which primarily target adaptive immunity. To date, no FDA-approved immunotherapies directly activate the innate anti-cancer immunity—an essential driver of lymphocyte recruitment and potentiator of responses to existing cancer immunotherapies. ENPP1, the dominant hydrolase that degrades extracellular cGAMP and suppresses downstream STING-mediated innate immune signaling, has emerged as a promising therapeutic target. However, existing ENPP1 inhibitors have been optimized for prolonged systemic residence time rather than effective target inhibition within tumors. Here, we report the characterization of STF-1623, a highly potent ENPP1 inhibitor with an exceptionally long tumor residence time despite rapid systemic clearance, enabled by its high ENPP1 binding affinity and slow dissociation rate. We show that membrane-bound ENPP1 on tumor cells, not the abundant soluble ENPP1 in serum, drives tumor progression. Consequently, STF-1623 unleashes anti-tumor immunity and synergizes with ionizing radiation, anti-PD-L1 and anti-PD-1, and a DNA damaging agent to produce robust anti-tumor and anti-metastatic effects across multiple syngeneic mouse tumor models, all without detectable toxicity. Conceptually, this work establishes that a noncovalent small molecule inhibitor of ENPP1 with ultralong drug-target engagement offers a safe and precise strategy to activate STING within tumors, fulfilling an unmet need of innate immunotherapies in cancer.

**One Sentence Summary:** A small molecule blocks ENPP1, reviving immune attack on tumors and enhancing immune therapy with minimal side effects in preclinical cancer models.

## INTRODUCTION

Despite the advancement in early detection and treatment, currently more than 18 million Americans are affected by cancer (*1*). Adaptative immune checkpoint inhibitors (ICIs) that block checkpoint proteins such as programmed cell death protein 1 (PD1), programmed death-ligand 1 (PD-L1), and cytotoxic T-lymphocyte associated protein 4 (CTLA-4) represent breakthroughs in cancer therapies in the past two decades. Adaptive ICIs unleash tumor infiltrating lymphocytes (TILs) to fight cancer, offering cures to patients who have exhausted other treatment options. Between 2015 and 2023, 11 ICIs were approved for over 20 general tumor types (*2*). However, only 20.13% patients are estimated to respond to ICIs as of 2023 (*2*). While tumors have a variety of strategies to develop ICI resistance, one of the key mechanisms is the lack of sufficient TILs in the first place for these ICIs to act on (*3,4*). Turning immunologically “cold” tumor “hot” by activating innate immune recognition of cancer cells and subsequently recruiting TILs has been the objective of the immune-oncology field for the past decade (*5*).

The stimulator of interferon genes (STING) pathway is a key innate immune pathway involved in anticancer immunity (*6–9*). STING is naturally activated by the second messenger 2′3′-cyclic-GMP-AMP (cGAMP), which is synthesized by the cytosolic double-stranded DNA (dsDNA) sensor cyclic-GMP-AMP synthase (cGAS) (*10–13*). Upon STING activation, the transcription factors interferon regulatory factor 3 (IRF3) and translocate to the nucleus to initiate the production of type-I interferons (IFN) (14,15). IFN has potent antitumor properties, and STING signaling through IRF3 is necessary for spontaneous clearance of immunogenic tumors in mice (*7,16*). In addition, STING also elicit antitumor immunity through IRF3-independnent pathways (*7,8*).

Given the importance of STING signaling in anticancer immunity, several cGAMP analogs have already entered clinical trials with reported results (NCT03937141; NCT03172936; NCT02675439; NCT04109092; NCT04096638; NCT06021626; NCT06626633), all of which pointing to limited efficacy with narrow therapeutic window (*17,18*). In addition to systemic inflammation, the disappointing result is likely due to the multifaceted roles that the STING pathway plays in different cell types within the tumor microenvironment (TME): although STING activation in dendritic cells and macrophages is responsible for eliciting antitumor immunity (*6,16,19*), STING activation in cancer cells paradoxically promotes metastasis (*20,21*). Additionally, the degree of STING activation can lead to different outcomes: while moderate STING activation is important for tumor vasculature normalization (*22*) and T cell function (*16,23*), extensive and prolonged STING activation causes vasculature toxicity in humans (*24*) and T cell deaths (*25–28*). This level of complexity highlights the need for more targeted approaches and offers insights into the limited clinical utility of STING agonists so far. However, aiming to achieve tumor specificity can also be problematic. For example, an antibody drug conjugate linking human epidermal growth factor receptor 2 to a direct STING agonist was created but resulted in one fatal event in phase 1 trial (NCT05514717), underscoring the power and danger of direct STING activation in the wrong cell types.

Instead of directly activating STING, we propose to target the dominant negative regulator of the STING pathway, ENPP1 (*29*), taking inspiration from adaptive ICIs that inhibit the negative regulators of the pathways. A hallmark of cancer is dsDNA mis-localization into the cytosol due to genomic instability (*21*), micronuclei (*30*), chromatin bridge (*31*), or extrachromosomal DNA (*32*). As a result, cancer cells with intact dsDNA sensor continuously produce cGAMP. To avoid self-STING activation-mediated cell death, cancer cells often suppress STING pathway through epigenetic silencing (*33*) or rewiring its downstream to noncanonical, pro-metastatic nuclear factor-kappa-B (NF-κB) pathway (*21*). Furthermore, cancer cells rapidly pump cGAMP out to the extracellular space for degradation by its extracellular hydrolase ENPP1 as another mechanism for immune evasion (*29,34,35*). ENPP1 is overexpressed by 50% solid tumors (*35*) and can be induced in non-cancerous bystander cells in the tumor TME including cancer-associated fibroblasts and macrophages (*34*). ENPP1 levels in tumor immune and stromal cells anti-correlate with their interferon signaling strength, suggesting that ENPP1 is a gatekeeper for cGAMP entry (*34*). *Enpp1* knockout in cancer cells or host unleashes the paracrine cGAS-STING signaling between cancer cells and immune cells utilizing cell-specific cGAMP importers (*26,36–41*), which delays tumor progression and abolishes metastases (*34*). Finally, outside of its role in downregulating the anti-cancer innate immune STING pathway, ENPP1 has also been shown to suppress tumor immunity by coupling with CD73 to generate immunosuppressive adenosine (*42,43*). Given these pro-tumor roles of ENPP1, we nominate ENPP1 as an ideal innate immune checkpoint target.

We previously reported a potent ENPP1 inhibitor STF-1623 (IC_50_ = 0.6 nM for human ENPP1, IC_50_ = 0.4 nM for mouse ENPP1) (*44*). However, STF-1623 exhibited fast serum pharmacokinetics in mice when administered systemically: its serum half-life was only 10-15 min, and the concentration dropped below IC_95_ (40 nM or 14.05 ng/mL) after 8 hours (*44*). However, in this study we found that systemically administered STF-1623 exhibits ENPP1 expression-driven tumor-selective targeting in mice. Co-crystal structure reveals structural determinants of STF-1623’s superior potency and selectivity towards ENPP1, giving rise to its long tumor residence time despite fast systemic clearance. We found that tumor-associated ENPP1, not the abundant serum ENPP1, plays a dominant role in tumor immune evasion, making STF-1623 a highly targeted and well-tolerated inhibitor. STF-1623 increased tumor cGAMP levels, induced tumor and serum IFN-γ production, and suppressed tumor growth and metastases with durable anti-tumor immunity across various tumor types. STF-1623 exploits cancer-produced extracellular cGAMP for controlled local activation of STING, essentially acting as a tumor-specific STING agonist.

## RESULTS

### Systemically administered STF-1623 concentrates in ENPP1 expressing tumors

To evaluate the pharmacokinetics of STF-1623 (**Fig. 1A**), we injected STF-1623 subcutaneously into BALB/c mice with established subcutaneous EMT6 breast tumors and measured the change in STF-1623 concentration with time. Although STF-1623 exhibited suboptimal pharmacokinetic properties (C_max_ = 23 µg/mL, t_1/2_ =1.7 hours) in the serum, it displayed much better pharmacokinetic properties in the EMT6 tumor (C_max_ = 9 µg/g, t_1/2_ = 6.6 hours) (**Fig. 1B**). In contrast, serum and tumor PK from mice bearing subcutaneous MC38 tumor were similar, both displaying fast half-times (t_1/2_ = 2.6 and 2.3 hours). (**Fig. 1C**). EMT6 and MC38 cancer cells differ significantly in their *Enpp1* expression (**Fig. 1D**) (*45*). We hypothesized that STF-1623 preferentially localizes to ENPP1-high EMT6 tumors due to interaction with its target. To test this hypothesis, we measured tumor and serum PK in mice bearing orthotopic WT or *Enpp1^-/-^* 4T1 tumors. Indeed, ENPP1 expression in cancer cells raised STF-1623 concentration and half-time (t_1/2_ = 4.3 vs. 1.1 hour) in the tumor (**Fig. S1A**). Additionally, as ENPP1 is ubiquitously expressed, we also assessed pharmacokinetics from tissues with the highest ENPP1 expression in healthy WT and *Enpp1-/-* BALB/c mice. We observed ENPP1-dependent increase in STF-1623 residence time in serum and liver, but not kidney, suggesting the latter as a main site of excretion (**Fig. S1B**). Together, STF-1623 exhibits target-driven tumor-selective localization, with ENPP1 expressed by cancer and bystander host cells both contributing to tumor retention. Long drug-target residence time (τ) is often observed in antibody drugs with slow dissociation kinetics from its target. To see if STF-1623 has slow dissociation from ENPP1, we quantified the kinetics and binding affinity between STF-1623 and ENPP1 using surface plasma resonance. Indeed, STF-1623 dissociate from mouse ENPP1 (K_off_ = (1.95 ± 0.58) x 10^-3^ s^-1^; τ = 540 ± 138 s) and human ENPP1 (K_off_ = (1.97 ± 0.41) x 10^-3^ s^-1^; τ = 524 ± 100 s) with exceptionally slow dissociation (K_off_) corresponding to ultralong τ, exhibiting a typical antibody-like binding profile (**Fig. 1E**).

**Fig 1.**
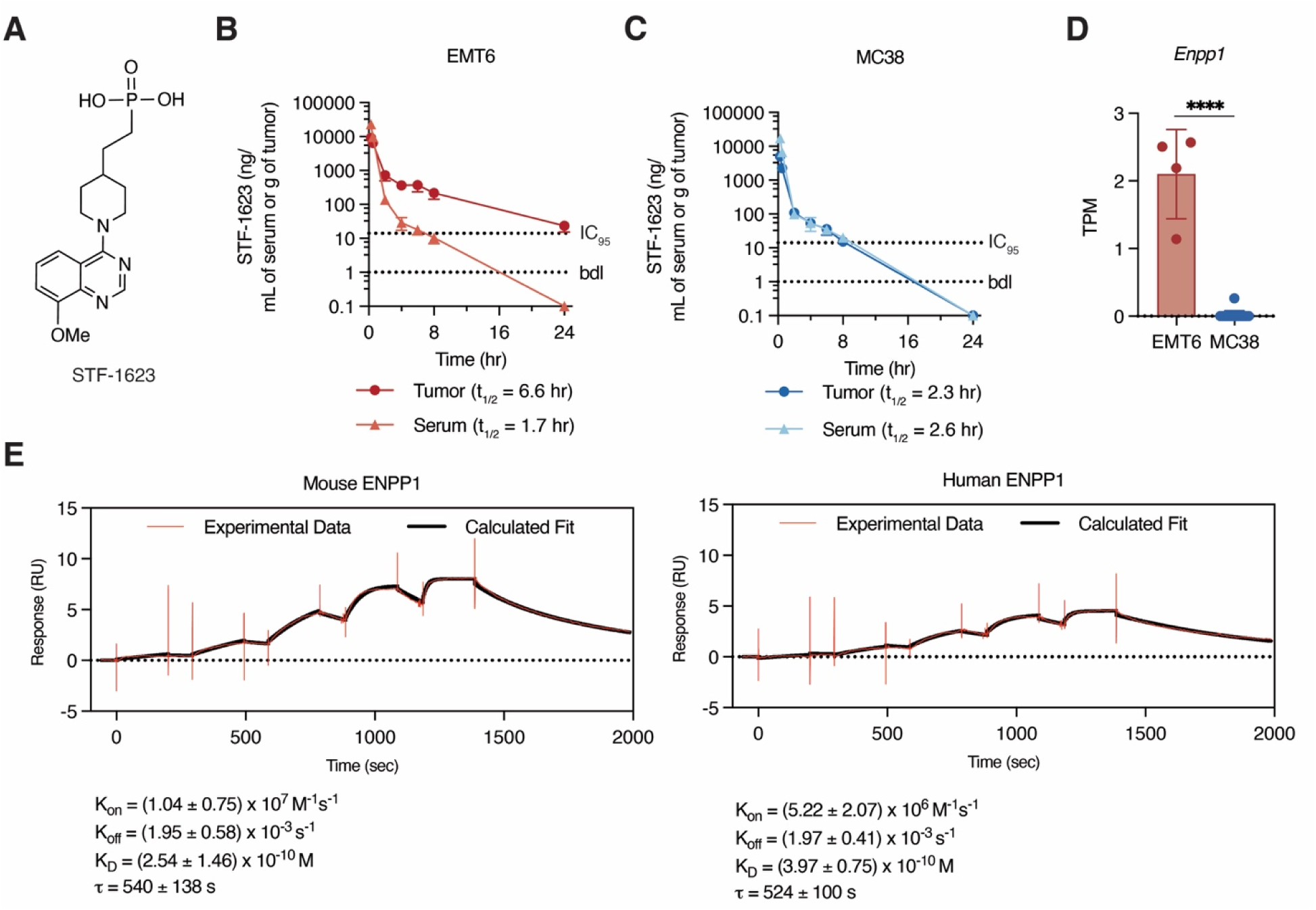
Systemically administered STF-1623 concentrates in ENPP1 expressing tumors. **(A)** Chemical structure of STF-1623. (**B**) Concentration of STF-1623 in serum and tumor of mice with established subcutaneous EMT6 tumors after one subcutaneous injection of STF-1623 (50 mg/kg). IC_95_ = 14 ng/mL or g; below detection limit (bdl) = 1 ng/mL or g. Mean ± SEM is plotted, n = 3 mice for each point except the following, where n = 2: tumor and serum 8 h (point removed as an outlier). (**C**) Concentration of STF-1623 in serum and tumor of mice with established subcutaneous MC38 tumors after one subcutaneous injection of STF-1623 (50 mg/kg. IC_95_ = 14 ng/mL or g; below detection limit (bdl) = 1 ng/mL or g. Mean ± SEM is plotted, n = 3 mice for each point except the following, where n = 2: serum 2 h (only 2 mice were collected); serum 4 h and 5 h (point removed as an outlier). (**D)** *Enpp1* RNA expression in EMT6 and MC38 cell lines. Mean ± SD is plotted. *P* value is determined by unpaired two-sided *t* test. TPM, transcript per million. **(E)** Single cycle kinetics analysis with surface plasmon resonance of the direct binding of 0.12, 0.36, 1.11, 3.33, 10 nM to N-terminally Avi-tagged and biotinylated mouse (left) or human (right) ENPP1 immobilized on a SA chip. Representative curves are plotted, n = 3 for moues ENPP1 and n = 4 for human ENPP1. Drug-target residence time (τ) is calculated as reciprocal of the dissociation rate constant (K_off_).\*\*\*\**P* < 0.0001.

### Co-crystal structure reveals molecular determinants of STF-1623’s potency and specificity towards ENPP1

The molecular determinants of STF-1623’s potency and long target engagement were unknown. Since crystallization attempts of human ENPP1 with STF-1623 yielded crystals that diffracted to low resolution, instead we created a proxy that mimics the catalytic site of ENPP1 in its paralog, ENPP3, which can readily yield high-quality crystals. ENPP3 shares identical amino acids within ∼4 Å of active site ligand with ENPP1 except for two residues (**Fig. 2A**). Therefore, we replaced those two residues, Q244 (which is K in ENPP1) and E275 (which is D in ENPP1), with the corresponding ENPP1 residues to create a faux ENPP1, hereby abbreviated as fxENPP1. Biochemically, fxENPP1 is a faithful proxy for ENPP1 (**Fig. 2B**): against native ENPP3, STF-1623 inhibits at an IC_50_ of 800 nM, over 1,000-fold lower potency compared to native ENPP1. Against fxENPP1, STF-1623 has an IC_50_ of 1.4 nM, indistinguishable from native ENPP1. We then solved the 2.7 Å co-crystal structure of fxENPP1 bound to STF-1623 using X-ray crystallography (**Fig. 2C,D**). Structural alignment with ENPP1 (PDB: 6WFJ) validated that the binding pocket is identical between the two proteins, further confirming that fxENPP1 is a faithful proxy for ENPP1(**Fig. 2E**). STF-1623 binds to the active site and forms extensive interactions with the fxENPP1 (**Fig. 2F**). The phosphonate head group binds the zinc that is essential for catalytic activity and also interacts with N226 (**Fig. 2F,G**). The piperidine linker forms hydrophobic interactions with L239 (**Fig. 2F,G**). The quinazoline group stacks between Y289 and F206 to form T[ - T[ interactions with both (**Fig. 2F,G**). Finally, D275 and K244 formed a hydrogen bonding network and perfect shape complimentarily with the 8-methoxy tail clamping down on the end of the compound (**Fig 2F,G**). These specific interactions could be responsible for the over 1,000-fold differences in potency of STF-1623 against human ENPP1 and human ENPP3 (**Fig. 2B**). Notably, the binding position of STF-1623 differs from that of STF-1084, an inhibitor with the same scaffold but different methoxy substituent positions, that we previously crystallized with mouse ENPP1 (*44*). Specifically, the nitrogens on the quinazoline of STF-1084 are pointed towards the hydrophobic back of the pocket and the 6,7-methoxys solvent-exposed, whereas STF-1623 nitrogens are solvent exposed and the 8-methoxy lies further back, a 180° flip accommodated by the piperidine linker (**Fig 2H**). In summary, STF-1623 is a potent and specific inhibitor of ENPP1 owing to the network of interactions it forms with ENPP1 enzymatic pocket.

**Fig 2.**
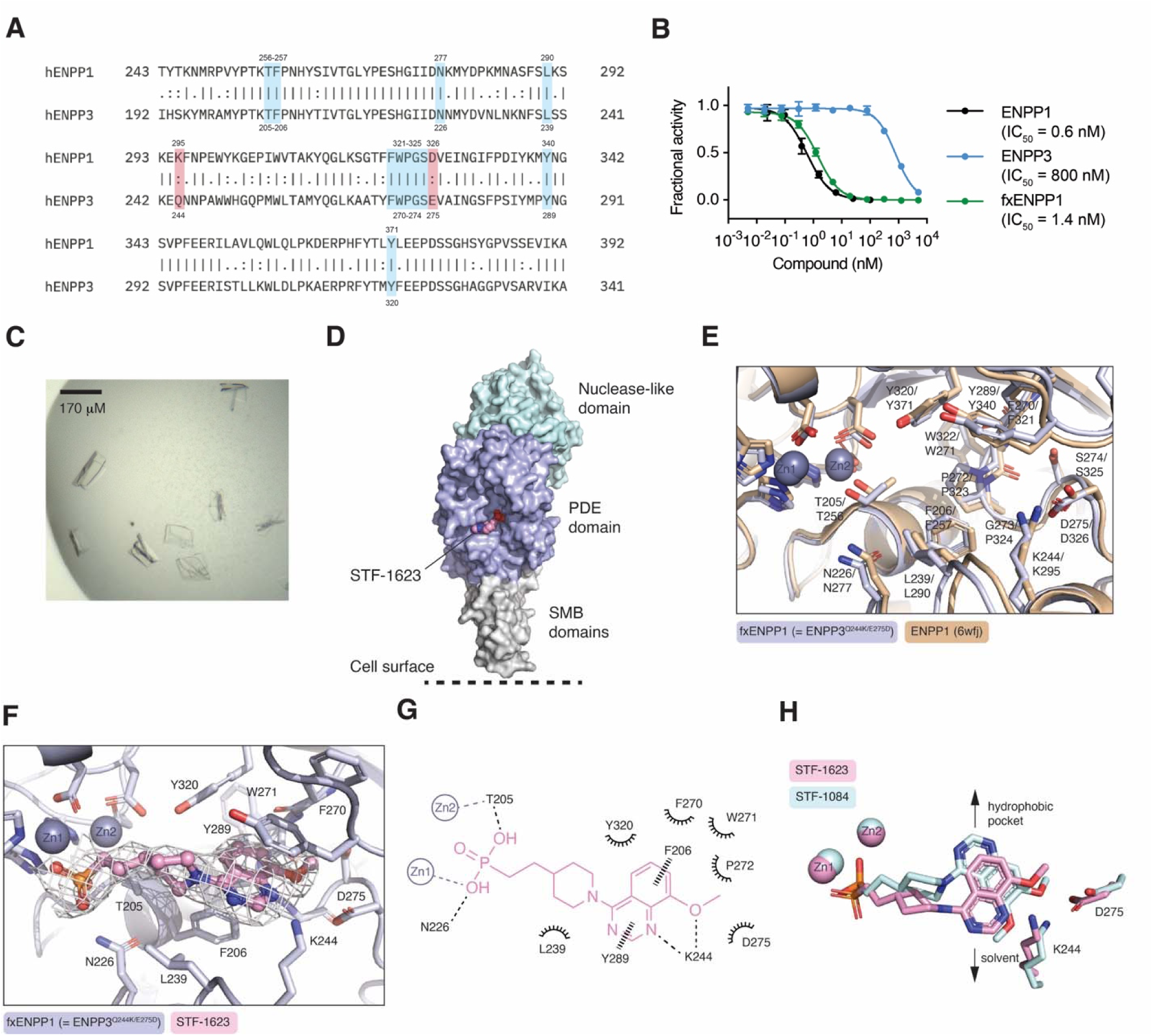
Co-crystal structure reveals molecular determinants of STF-1623’s potency and specificity towards ENPP1. (**A**) Sequence alignment between human ENPP1 and ENPP3. Residues within 4Å of the active site are highlighted in blue if they are the same between human ENPP1 and ENPP3, and in pink of they are different. (**B)** *In vitro* dose-inhibition curves to determine IC_50_ values of STF-1623 against purified human ENPP proteins. Enzyme concentrations of 15 pM (ENPP1) or 250 pM (ENPP3 and fxENPP1) and cGAMP concentration of 2 μM were used. cGAMP degradation was measured with cGAMP-luc assay. Dots represent the mean of 2 biological replicates. (**C**) Image of the crystal. (**D**) The 2.7 Å crystal structure of STF-1623 (pink spheres) bound to human fxENPP1 (teal surface: nuclease-like domain; purple surface: phosphodiesterase domain; gray surface: somatomedin B domain). (**E**) Structural alignment between fxENPP1 (=ENPP3^Q244K/E275D^, purple) and ENPP1 (wheat, PDB: 6wfj). (**F**) Expanded view of STF-1623 (pink sticks/spheres) bound in the active site of fxENPP1 (purple sticks/cartoon). Zincs are shown as dark purple spheres. Electron density of STF-1623 shown as gray mesh, 0.7 rmsd. (**G**) Schematic drawing of interactions formed between STF-1623 (pink) and the fxENPP1 active site residues (black). Residues within 4Å are shown (not including the zinc-coordinating residues). Metal coordination shown as gray dashed lines, hydrogen bonds shown as black dash lines, aromatic interactions shown as black wedged lines, and hydrophobic or polar interactions shown as spokes. (**H**) Overlay of STF-1623 (pink) with STF-1084 (blue, PDB: 6XKD), bound to human fxENPP1 and mouse ENPP1, respectively. Ligands are shown as sticks. Protein residues K244, D275, and zincs are shown.

### Intratumoral ENPP1 membrane retention determines tumor progression

In addition to being a transmembrane protein, ENPP1 is also secreted (**Fig. 3A**) and is detected at high levels (2.8 µg/L) in serum (The Human Protein Atlas). We previously showed high potency of STF-1623 at inhibiting serum ENPP1 activity (*44*), suggesting it also binds to secreted ENPP1 (secENPP1). SecENPP1 has a quick clearance rate (*46*) that likely contributes to STF-1623’s fast clearance rate. Additionally, the inhibitor is highly hydrophilic (*44*). Any compound not bound to ENPP1 would likely remain unbound to other serum protein and undergo fast excretion by the kidney (*44*). Given the different pharmacokinetic properties of STF-1623 in the tumor and serum targeting membrane ENPP1 (memENPP1) and secENPP1, respectively, it is important to which of the two isoforms are the active target. Despite the differences in disulfide-mediated dimerization (**Fig. S2A,B**), both memENPP1 and secENPP1 are active towards cGAMP (**Fig. 3B**) and ATP (**Fig. S2C**). Using purified enzymes (**Fig. S2D**), we determined that memENPP1 (*K_cat_* = 189 s^-1^, *K_m_* = 313 µM, *K_cat_/K_m_* = 6.1 x 10^5^ M^-1^ s^-1^) and secENPP1 (*K_cat_* = 54 s^-1^, *K_m_* = 101 µM, *K_cat_/K_m_* = 5.5 x 10 M s) both efficiently hydrolyze cGAMP (**Fig. 3C**).

**Fig 3.**
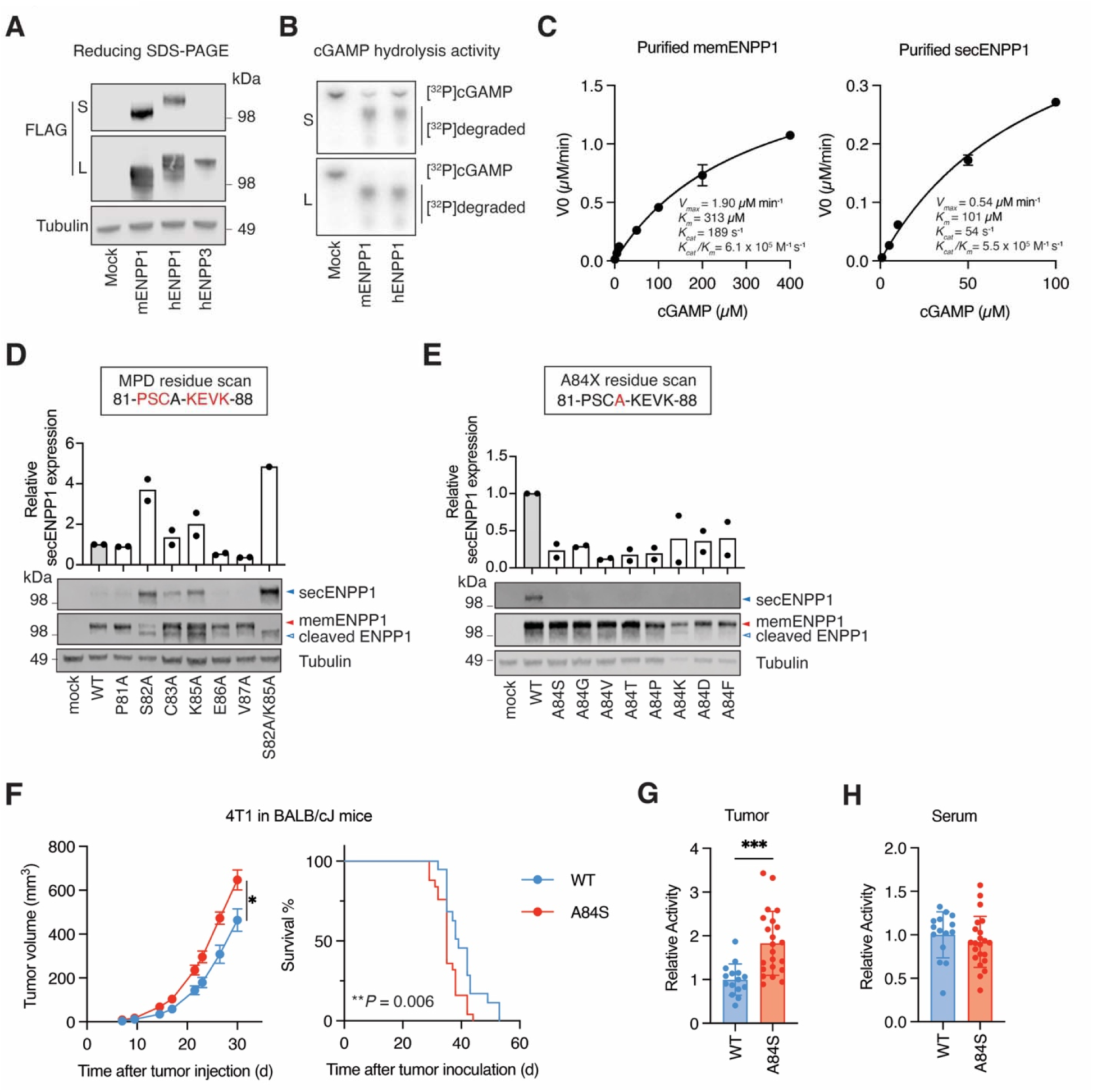
Intratutmoral ENPP1 membrane retention determines tumor progression. (**A**) Expression of mENPP1, hENPP1, hENPP3 in supernatant and lysate of 293T *ENPP1^-/-^*cells assessed by reducing western blotting. Data are from one experiment (full scan of blot available as source data). (**B**) Activity of mENPP1 and hENPP1 in supernatant and lysate assessed by [^32^P] cGAMP hydrolysis by thin-layer chromatography after 4 hours of reaction at pH 9.0. Data are from one experiment. (**C**) Kinetics of cGAMP hydrolysis by 10 nM purified memENPP1 and secENPP1 at pH 9.0. Mean ± SEM is plotted, n = 2 independent experiments for memENPP1; n = 3 technical replicates from 2 independent experiments for secENPP1. Data are fit with Michaelis-Menten model. (**D**) and (**E**), Expression of WT ENPP1, MPD residue mutants (D), and A84 residue mutants (E) in supernatant and lysate of 293T cGAS *ENPP1^-/-^* cells assessed by reducing western blotting. memENPP1, secENPP1, and cleaved ENPP1 are indicated as red, blue closed, and blue open triangles respectively. Relative secENPP1 expression is calculated as secENPP1 over the sum of secENPP1 and memENPP1. Quantification is from two independent experiment, with blots from one representative experiment shown (Full scan of blot available as source data). (**F**) 4T1^WT-OE^ (*n* = 20 mice from two independent experiments), 4T1^A84S-OE^ (*n* = 25 mice from two independent experiments) cells (2.5 or 5 x 10^4^) were orthotopically injected in WT BALB/c mice. One mouse without tumor engraftment was excluded. Mean ± SEM is plotted. The *P* value for tumor volume on day 30 was determined by multiple unpaired *t* test, while for Kaplan-Meier curve was determined by log-rank Mantel-Cox test. (**G**) and (**H**), Relative cGAMP hydrolysis activity from tumor lysates (G) and sera (H) from randomly selected mice in (F) reaching experimental endpoints (n = 15, 22 for 4T1^WT-OE^ and 4T1^A84S-OE^, respectively). Mean ± SD is plotted. *P* value is determined by unpaired two-sided *t* test. mENPP1: mouse ENPP1; hENPP1: human ENPP1; hENPP3: human ENPP1; S: supernatant; L: lysate; memENPP1: transmembrane ENPP1; secENPP1: secreted ENPP1; MPD: membrane proximal domain. \**P* < 0.05, \*\**P* < 0.01, \*\*\**P* < 0.001.

Since both forms of the protein have similar activity, we needed an approach for selective perturbation. This led us to explore the mechanism of secretion. SecENPP1 is not produced from alternative splicing (*47*), ectodomain shedding (*48*), or alternative translation start site (**Fig. S3A**). Instead, secENPP1 could be proteolytically processed by the signal peptide complex given a predicted cleavage motif at A84/K85 (**Fig S3B,C**) (*49*). To test this, we first performed an alanine mutation scan around the putative cleavage site and observed increased secENPP1 expression in S82A and K85A (**Fig. 3D**). Next, we mutated A84 residue, -1 position from the putative cleavage site required to be occupied by small non-charged residues (*50*), to a variety of other amino acids; all compromised the production of secENPP1 (**Fig. 3E**). Overall, mutagenesis-induced changes in ENPP1 secretion correlated with the predicted secretion probability based on consensus signal peptide sequence, confirming the mechanism of secENPP1 production (**Fig. S3D**). Intriguingly, we also identified a mutant A84S that compromises secENPP1 production (**Fig. 3E**) that corresponds to a human single nucleotide polymorphism (rs125086092). This variant that alters the ratio of membrane bound to secreted ENPP1 provided a tool to dissect the functions of the two forms of ENPP1 *in vivo* (**Fig. S3E**).

Next, we sought to determine how 4T1 tumors expressing different ratios of mem:secENPP1 would influence protein localization and tumor growth. We observed that 4T1 cells overexpressing ENPP1 mutant that retains all ENPP1 on the plasma membrane (4T1^A84S-OE^) promoted tumor growth faster than 4T1^WT-OE^ that partially secretes its ENPP1 (**Fig. 3F**). We collected tumors and sera from these mice at a humane endpoint. ENPP1^A84S-OE^ cells with increased surface tethering of ENPP1 led to increased intratumoral ENPP1 activity (**Fig. 3G**). Conversely, ENPP1^WT-OE^ tumors that shed secENPP1 had slightly elevated serum ENPP1 activity, suggesting that secENPP1 from tumors is potentially cleared into the circulation (**Fig. 3H**). To formally test this, we measured serum cGAMP activity from mice bearing 4T1^WT-OE^ tumors of various sizes and observed a size-dependent increase in serum cGAMP activity (**Fig. S3F**), a trend that disappeared in mice bearing 4T1^A84S-OE^ that does not shed its ENPP1 (**Fig. S3G**). These data confirmed that while memENPP1 on cancer cells is retained within the tumor, secENPP1 are cleared into circulation. Since tumor growth correlated with intratumoral but not serum cGAMP degradation activity, this suggests that memENPP1 exerts regulation of extracellular cGAMP locally in the tumor, which subsequently influences tumor progression. Together, we delineated the role of memENPP1 in tumor progression and propose that ENPP1 blockade therapies need to target tumor ENPP1. STF-1623 preferentially targets ENPP1-high tumors, fulfilling this criterium.

### STF-1623 synergizes with anti-PD-L1 and ionizing radiation to abolish EMT6 breast cancer metastasis

With the optimized pharmacokinetic properties, we next characterized the pharmacodynamics of STF-1623 in the subcutaneous EMT6 breast tumor model. Using mass spectrometry, we found that homogenized EMT6 tumors treated with STF-1623 showed a time-dependent increase in tumor cGAMP (presumably both extracellular and intracellular) levels over vehicle treated mice with an approximate doubling within hours and lasting for over 24 hours (**Fig. 4A**). cGAMP was not detected in the serum (below low limit of quantification of 2 ng/mL) across all samples independent of treatment group. We also observed an increase in interferon gamma (*Ifn-*γ) mRNA levels in tumors as soon as 15 minutes (**Fig. 4B**) and an elevated serum IFN-γ concentration to ∼ 5 pg/mL after 24 hours (**Fig. 4C**), all of which demonstrate target inhibition in the tumors. Conversely, MC38 tumors without intratumoral STF-1623 accumulation lacked cGAMP-STING mediated PD effects as tumor cGAMP or serum IFN-γ levels were not increased (**Fig. S4A,B**).

**Fig 4.**
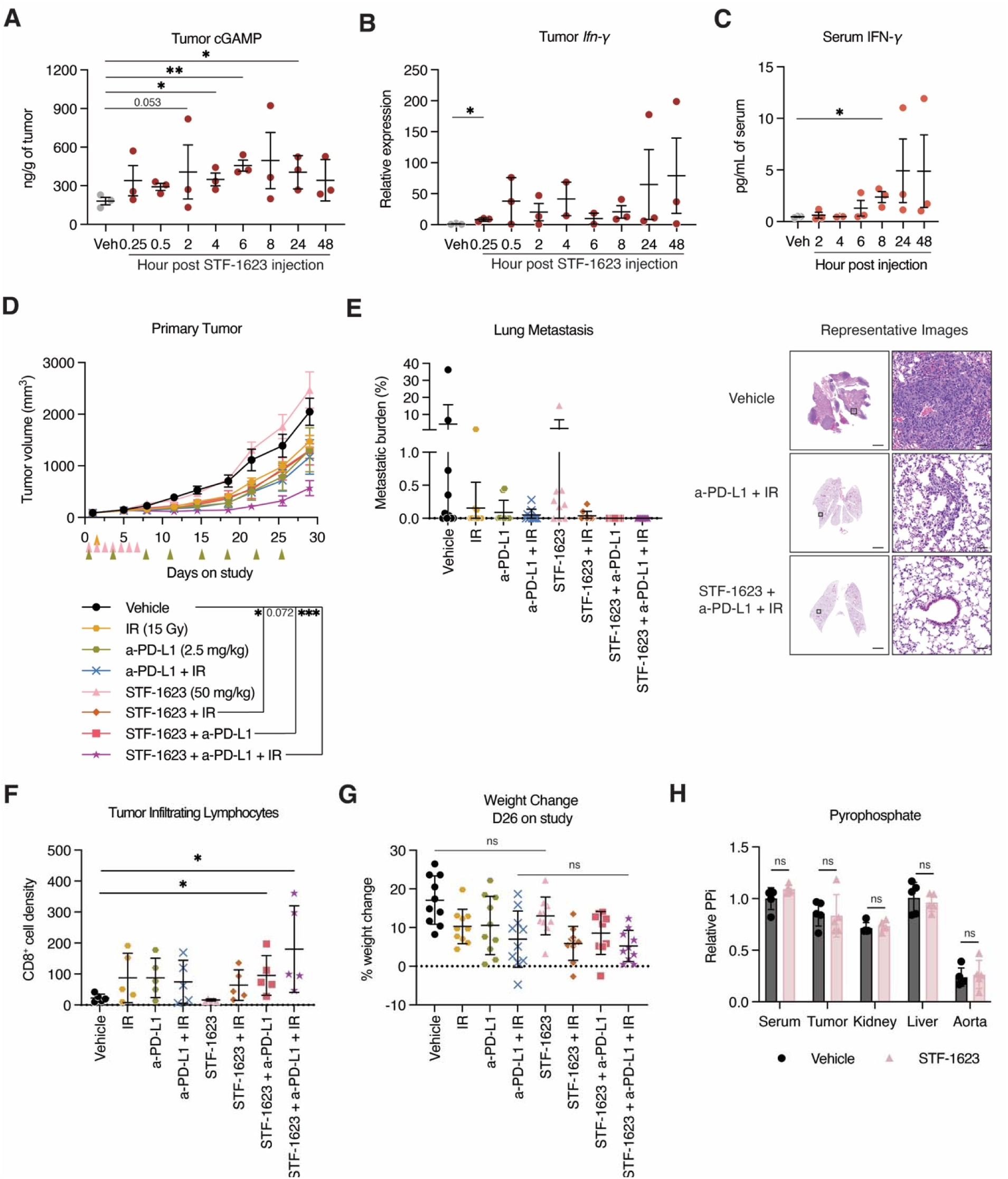
STF-1623 synergizes with a-PD-L1 and ionizing radiation to abolish EMT6 breast cancer metastasis. (**A**) cGAMP amount (ng/g) in subcutaneous EMT6 tumors at time points indicated after one dose of STF-1623 (50 mg/kg) subcutaneous injection. Mean ± SEM is plotted, n = 3 mice. (**B**) Relative *Ifn-γ* mRNA expression in subcutaneous EMT6 tumors at time points indicated after one dose STF-1623 (50 mg/kg) subcutaneous injection. Mean ± SEM is plotted, n = 2-3 mice (average of technical triplicates). (**C**) IFN-γ levels (pg/mL) in serum of EMT6 bearing mice at time points indicated one dose STF-1623 (50 mg/kg) subcutaneous injection. Mean ± SEM is plotted, n = 3 mice (average of technical duplicates). (**D**) EMT6 cells (5 x 10^5^) were subcutaneously injected in BALB/c mice. Mice with established tumors were randomized into 10 per group and received ionizing radiation (IR), anti-PD-L1 (a-PD-L1), STF-1623 or their combinations at the dosage and frequencies (orange, green, pink triangles respectively) indicated. Mean ± SEM is plotted. The *P* value for tumor volume on day 29 was determined by multiple unpaired *t* test. (**E**) Percent lung metastatic burden determined by hematoxylin and eosin (H&E) staining of mice in (D) terminated on day 29 of the study. Mean ± SD is plotted, n = 9-10 mice. Representative H&E images of the whole lungs (left, scale bar is 4 mm) and zoomed in areas (right, scale bar is 0.1 mm) are shown. (**F**) Immunohistochemistry of CD8^+^ T cells in randomly selected tumors of mice in (D) terminated on day 29 of the study. Mean ± SD is plotted, n = 5 mice (representative images shown in Fig. S5B). (**G**) Percent weight change of mice in (D) on day 25 compared to day 1 of the study. Mean ± SD is plotted, n = 9-10 mice. (**H**) Relative pyrophosphate (PPi) levels in serum, orthotopic breast tumor, kidney, liver, aorta of mice treated with vehicle or STF-1623 (50 mg/mL) subcutaneously daily for seven days. Mean ± SD is plotted, n = 5 mice (average of two technical replicates). *P* values were determined by two-sided unpaired *t* test unless otherwise mentioned. \**P* < 0.05, \*\**P* < 0.01, \*\*\**P* < 0.001; P value is shown if between 0.05 and 0.1; not significant (ns).

We proceeded to test the anti-tumor efficacy of STF-1623 as a monotherapy and in combination with anti-PD-L1 (a-PD-L1) or ionizing radiation (IR) at optimized dose of 15 Gy (**Fig. S5A**). While STF-1623 did not exhibit single agent efficacy in EMT6 tumors, it synergized with a-PD-L1 and IR to shrink primary tumors and abolish lung metastases (mice with lung metastasis: vehicle 6/10 vs. STF-1623 + a-PD-L1 + IR 0/10, *P* = 0.0108) (**Fig. 4D**). Additionally, STF-1623 with a-PD-L1 dual combination therapy also completely abolished lung metastases (vehicle 6/10 vs. STF-1623 + a-PD-L1 0/9, *P* = 0.0108) (**Fig. 4D**). This result agrees with our previous observation of the deterministic role that ENPP1 status plays in predicting long-term metastasis of breast cancer patients receiving a-PD-1 neoadjuvant therapy (*34*). STF-1623 + a-PD-L1 dual and STF-1623 + a-PD-L1 + IR triple combinations both lead to significant increase in CD8^+^ tumor infiltrating lymphocytes by immunohistochemistry (**Fig. 4F**), supporting that STF-1623 acts through immunomodulation. In contrast to EMT6, the MC38 tumors that did not exhibit the desired pharmacokinetic and pharmacodynamic profiles was not affected by STF-1623 alone or in combination with anti-PD-1 (a-PD-1) (**Fig. 4C,D**). Rather, the immunologically “hot” MC38 tumors (*51*) were sensitive to a-PD-1 treatment alone (**Fig. S4C**).

STF-1623 alone or in combination therapy is well-tolerated as no significant weight loss was observed across groups (**Fig. 4G**). In terms of on-target side-effects, besides cGAMP, ENPP1 also degrades extracellular adenosine triphosphate (ATP) to pyrophosphate (PPi), which is critical to calcium homeostasis (*52,53*). Previously, we demonstrated that tissue ENPP1 ATP hydrolysis activity, hence tissue PPi level, is necessary and sufficient for maintaining calcium homeostasis (*52*). We measured PPi levels from breast cancer bearing mice after seven-day dosing of STF-1623 (50 mg/kg) versus vehicle controls and found no significant differences in PPi levels in serum, tumor, and organs where ENPP1 expression is known to be high and/or dysregulation of calcium homeostasis could result in pronounced pathology (**Fig. 4H**). This is mostly likely due to the redundant enzymatic function of ENPP3 (*54*), short course of ENPP1 treatment, and quick clearance of STF-1623 from the circulation. Together, STF-1623 is safe, has minimal on-target and off-target side effects, and exhibits anti-metastatic effects in murine breast cancer.

### STF-1623 controls Panc02 pancreatic and CT26 colorectal tumors by activating anti-cancer immunity

Next, we investigated STF-1623’s efficacy in other cancer types. We have previously shown the tumor shrinkage effect of STF-1623 as a single agent and in combination with IR in the Panc02 mouse pancreatic tumor model, when infused continuously through an osmotic pump for 7 days (*35*). The desirable tumor pharmacokinetics and pharmacodynamics of daily subcutaneous dosing of STF-1623 indicated that continuous infusion is not necessary. To optimize treatment dosage and duration of STF-1623 in Panc02, we performed flow cytometry on Panc02 tumors 12 days after treatment initiation. Overall, although STF-1623 monotherapy did not affect immune cell numbers and functions compared to vehicle control, STF-1623 and IR combination therapy resulted in pronounced increase in the ratio between pro-inflammatory, anti-cancer M1 (IA-IE^+^CD206^low^ F4/80^+^GR-1^-^) and anti-inflammatory, pro-cancer M2 macrophages (IA-IE^-^ CD206^high^F4/80^+^GR-1^-^) (M1/M2), the number of CD335^+^CD3^-^ natural killer (NK) cells, CD335^+^CD3^+^ NK T (NKT) cells, CD4^+^ T cells, CD8^+^ T cells, as well as the ratio between CD8^+^ T cells and Foxp3^+^ regulatory T (Treg) cells in the tumors (**Fig. 5A**). In terms of T cell function, STF-1623 and IR increased the number of activated (as shown by the early activation marker CD69 and late activation marker PD-1) and proliferating (Ki67^+^) CD4^+^ and CD8^+^ T cells compared to vehicle controls and IR alone (**Fig. 5B**). The immunostimulatory effects of STF-1623 is dose- and duration-dependent: only doses at or above 5 mg/kg and duration at or above 3 days reprogrammed the tumor microenvironment (**Fig. 5A,B**). We noticed a mild decrease in total and proliferating CD4^+^ T cells when increasing STF-1623 (5 mg/kg) from 3-day dosing to 7-day dosing, suggesting of potential CD4+ T cell killing by high levels cGAMP upon ENPP1 inhibition (**Fig. 5A,B**), consistent with our previous report (*25*).

**Fig 5.**
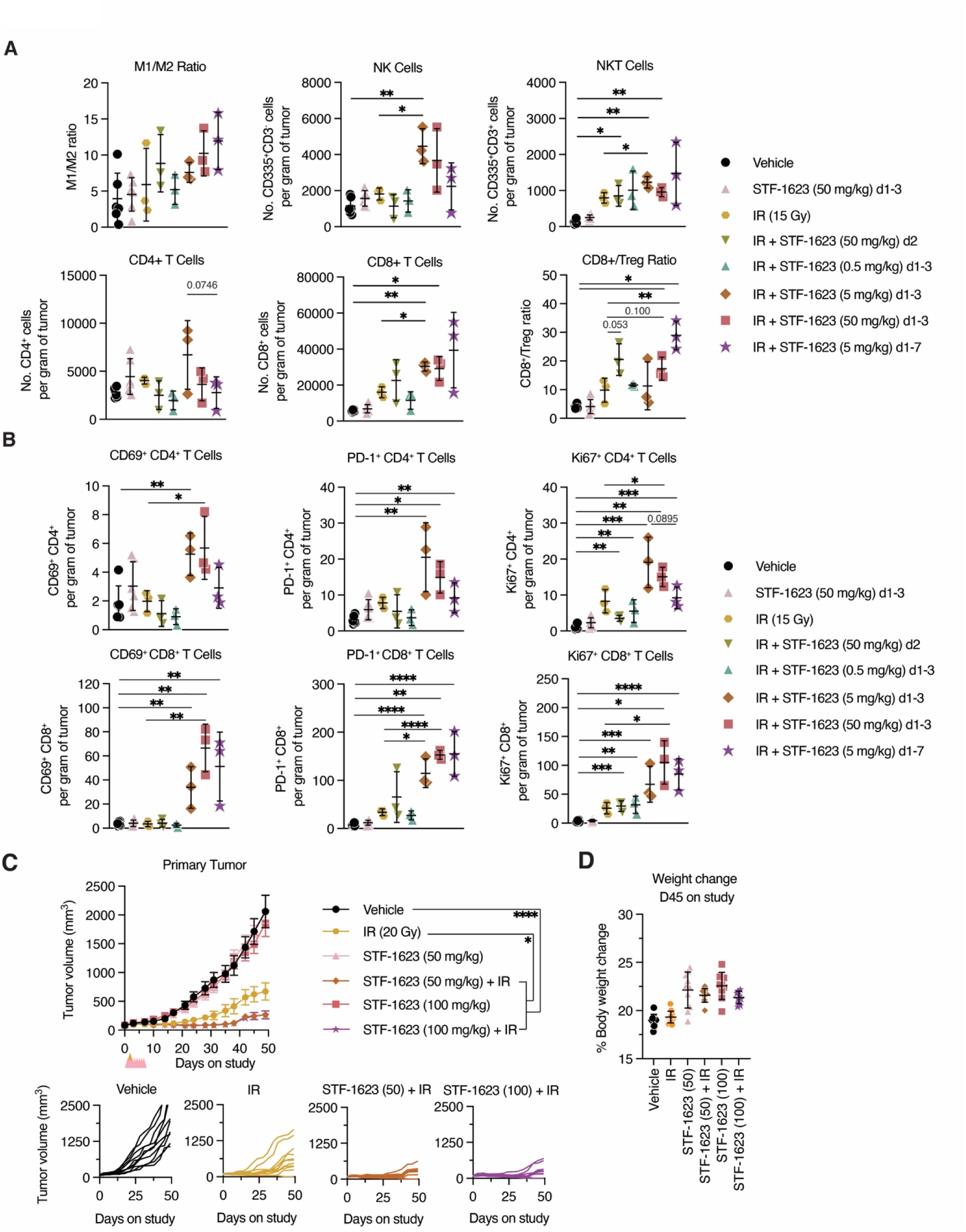
STF-1623 controls Panc02 pancreatic tumor growth by activating anti-cancer immunity. (**A**) and (**B**) Panc02 (3 x 10^6^) were subcutaneously injected into female BALB/c mice, randomized when tumors established, and received treatment indicated. On day 12 of the study (day 18 post tumor inoculation), tumors were isolated and processed for flow cytometry (raw gating available as source data). Mean ± SD is plotted, n = 6 mice for vehicle and STF-1623 groups, and n = 3 for all other combination therapy groups. The percentage of IA-IE^+^CD206^low^ M1 macrophage (F4/80^+^GR-1^-^) over IA-IE^-^CD206^high^ M2 macrophage, the number of CD335^+^CD3^-^ NK cells, CD335^+^CD3^+^ NKT cells, CD4^+^CD3^+^ T cells, CD8^+^CD3^+^ T cells per gram of tumors, and ratio of CD8^+^ T cells over Foxp3+CD4+ Treg cells (A), CD69^+^CD4^+^ T cells, PD-1^+^CD4^+^ T cells, Ki67^+^CD4^+^ T cells, CD69^+^CD8^+^ T cells, PD-1^+^CD8^+^ T cells, and Ki67^+^CD8^+^ T cells (B). (**C**) Panc02 (3 x 10^6^) were subcutaneously injected into female BALB/c mice, randomized when tumors established, and received STF-1623, IR, or their combinations at dosage and frequencies indicated (STF-1623: pink triangle, IR: orange triangle). Mean ± SEM is plotted, n = 9-12 mice per group. The *P* value for tumor volume on day 29 was determined by multiple unpaired *t* test. Spider plots of individual tumor growth are shown. (**D**) Percent weight change of mice in **c** on day 45 compared to day 0 of the study. Mean ± SD is plotted, n = 9-12 mice. NK: natural killer; NKT: natural killer T; Treg: regulatory T cells. IR: ionizing radiation. *P* values were calculated by two-sided unpaired *t* test unless otherwise noted. \**P* < 0.05, \*\**P* < 0.01, \*\*\**P* < 0.001; P value is shown if between 0.05 and 0.1.

To investigate if the immunostimulatory effects of STF-1623 in combination with IR could elicit tumor control, we delivered STF-1623 (50 mg/kg or 100 mg/kg) subcutaneously daily for seven days. Although STF-1623 alone did not affect Panc02 tumor growth, combination of STF-1623 at either dosage with IR completely suppressed tumor growth for over 30 days after treatment completion (**Fig. 5C**). The tumor growth inhibition effect of STF-1623 and IR combination therapy was significantly larger than no treatment or IR alone (**Fig. 5C**). Lastly, this treatment regimen is well-tolerated by the animals (**Fig. 5D**).

Similar results were observed in the CT26 colorectal tumors that responded to a-PD-1 (**Fig. S6A**) but less sensitively than MC38 colorectal tumors (**Fig. S4C**). STF-1623 not only showed comparable single agent efficacy as a-PD-1 but also acted synergistically with a-PD-1 to control tumor growth, while being well-tolerated (**Fig. S6A,B**). Moreover, we observed dramatic increase in tumor infiltrating CD4^+^ and CD8^+^ T cells in a-PD-1 combined with STF-1623 at dosage as low as 5 mg/kg (**Fig. S6C**). Together, our results demonstrated that STF-1623 synergizes with IR and a-PD-1 to control mouse pancreatic and colorectal cancer growth, respectively, through reprogramming an immunosuppressive TME to an immunostimulatory one.

### STF-1623 crosses the blood brain barrier and exhibits efficacy in delaying GL261 glioblastoma growth

ENPP1 is a potential target in glioblastoma, as ENPP1 is highly expressed in the brain (*54*), and the expression in glioblastoma (GBM) stem cells maintains its stem-like properties (*55*). Therefore, we investigated if STF-1623 (50 or 100 mg/kg) can cross the blood brain barrier (BBB) when dosed systemically. Once again, a serum t_1/2_ of less than 2 hours was observed with no detectable STF-1623 present at 24 hours (**Fig. 6A**). Surprisingly, STF-1623 was detected in both the cerebrospinal fluid (CSF) and the brain parenchyma within 5 min of drug dosing, and its level persisted above its IC_95_ even at 24 h, at which time it has been cleared from the serum, suggesting the measured amount is unlikely serum contamination (**Fig. 6A**). Comparison of AUC_0-inf_ values (serum: 18,333 ng•hr/ mL; CSF: 1,980 ng•hr/ mL; brain: 751 ng•hr/ mL STF-1623 when injected at 50 mg/kg) suggest ∼5-10% of STF-1623 crossed the BBB into CSF and brain parenchyma. These pharmacokinetics data suggest that to a limited degree, STF-1623 can cross an intact BBB and reside at low levels for an extended period in CSF and brain parenchyma. We next tested its efficacy in the murine GL261 Red-FLuc (GL261) GBM model established intracranially. The standard of care for GBM includes IR plus the alkylating agent temozolomide (TMZ). We reasoned that this combination would increase cGAMP production from mislocalized dsDNA thereby synergizing with STF-1623. Indeed, while TMZ alone and in combination with IR drastically increased survival, TMZ + IR + STF-1623 triple therapy exhibited superior efficacy compared to no treatment, TMZ alone, TMZ with IR or TMZ with STF-1623 (**Fig. 6B**). After 28 days of treatment initiation, 5/8 (62.5%) mice receiving triple combination therapy remained completely tumor free (**Fig. 6C**) while tolerating the regimen well (**Fig. 6D**). In summary, STF-1623 is a potent immune checkpoint blocker in various murine solid cancers including brain cancers.

**Fig 6.**
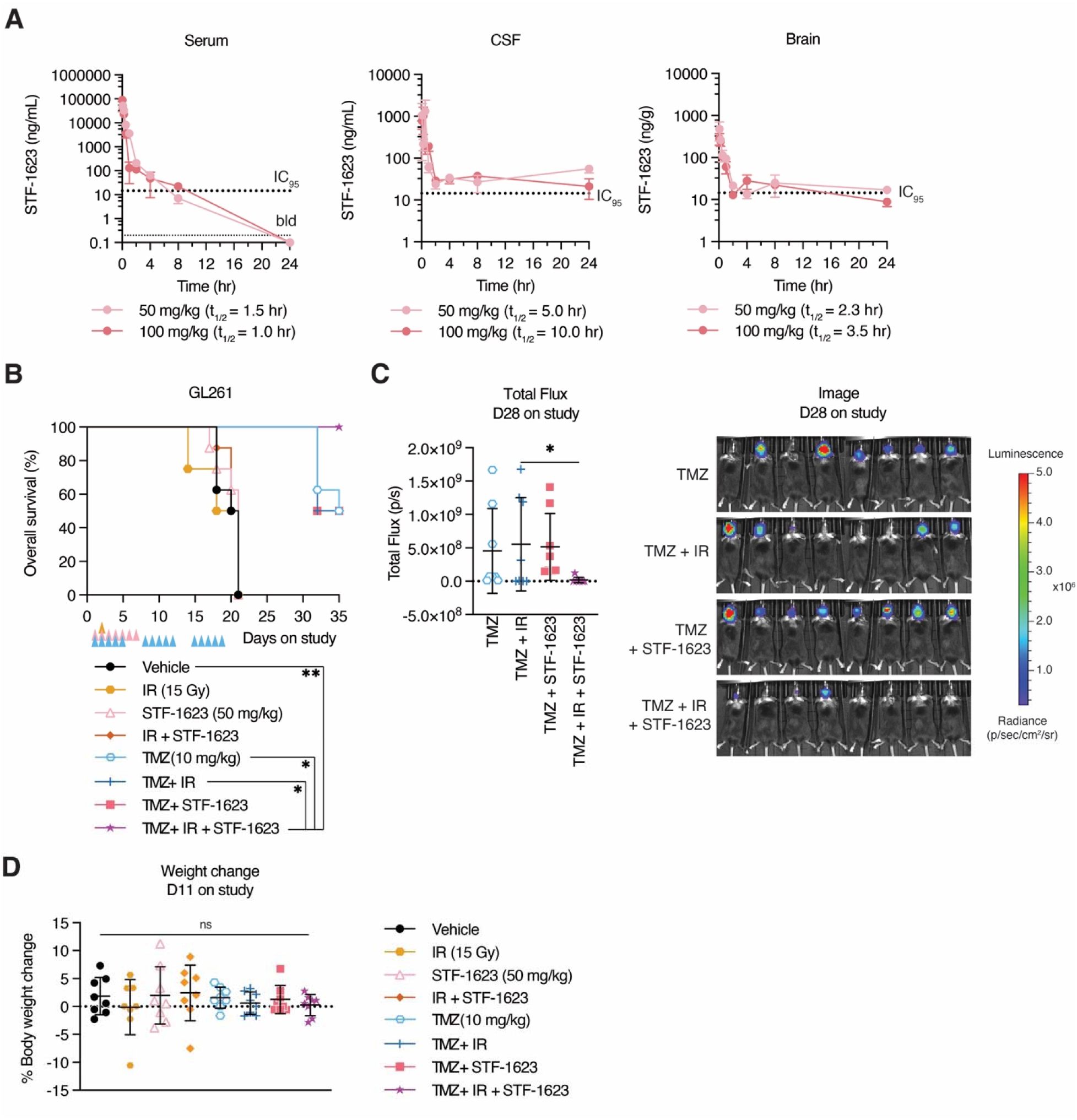
STF-1623 cross the blood brain barrier and exhibits efficacy in delaying GL-261 glioblastoma growth. (**A**) Concentration of STF-1623 in serum, cerebral spinal fluid (CSF), and brain of mice after one dose of STF-1623 (50 or 100 mg/kg) via subcutaneous injection. IC_95_ = 14 ng/mL or g; below detection limit (bdl) = 1 ng/mL or g. Mean ± SEM is plotted, n = 3 mice for each point except the following, where n = 2: serum (50 mg/kg) at 0.08 and 1 hour; serum (100 mg/kg) at 4 hours (points removed as outliers). (**B**) GL261 Red-Fluc (3 x 10^5^) were intracranially injected in C57BL/6 mice. Five days after tumor inoculation (day 1 of study), mice received a dose of IR, TMZ orally, STF-1632 subcutaneously or a combination of the three at dosage and frequency indicated (STF-1623: pink triangle, IR: orange triangle; TMZ: blue triangle). Kaplan-Meier of survival determined to death or humane end point is plotted, n = 8 mice. The *P* value was determined by determined by log-rank Mantel-Cox test. (**C**) Quantification and raw images of the total flux (p/s) of mice in (B) on day 28 of the study of groups with TMZ. Mean ± SD is plotted, n = 8 mice. (**D**) Percent weight change of mice in (B) on day 11 of the study. Mean ± SD is plotted, n = 9-10 mice. IR: ionizing radiation; TMZ: temozolomide. *P* values were calculated by two-sided unpaired *t* test unless otherwise noted. \**P* < 0.05. \*\**P* < 0.01.

## DISCUSSION

Cancer therapeutics have long faced the challenge of tumor specificity, and therefore intolerable side-effects which often prevents the use of curative dosage. Towards this end, drugs with long τ in tumors but fast systemic clearance have been sought after to ensure efficacy and safety. This is particularly relevant in targeting ENPP1, as although it exits in abundance in the circulation, it is the tumor-associated ENPP1 that dictates tumor progression, thus constituting the active target. Efforts have been dedicated to increasing the τ of small-molecule inhibitors in cancer therapy (*56*), including the development of covalent inhibitors (*57*). However, covalent inhibitors are frequently limited by the lack of available cysteines and other binding residues near the substrate binding site of the target, such as in the case of ENPP1. Here, we discovered that STF-1623, although a noncovalent inhibitor, has superior slow K_off_ owing to its high binding affinity, which gives rise to its long τ specifically in tumors. Ultralong τ allows for daily dosing regimen, which achieved efficacy without affecting ENPP1’s roles in calcium homeostasis in normal tissue. Another challenge of inhibiting ENPP1 is target specificity. STF-1623’s potency towards human ENPP1 is dictated by the identity of just two amino acids in the enzymatic pocket, K244 and D275, rendering it specific towards human ENPP1 but not its paralog ENPP3 (*54*). As ENPP1 and ENPP3 have different effects on cancer (*54*), this specific inhibitor enables precise targeting of ENPP1 while avoiding potential side-effects from dual ENPP1- and ENPP3-inhibition. Together, STF-1623 is a ENPP1-specific inhibitor that homes to tumors with long τ and fast systemic clearance. Noncovalent small-molecule inhibitors with ultralong τ like STF-1623 represents a more widely applicable strategy for drug development.

By comparing pharmacokinetic and pharmacodynamic properties of STF-1623 between ENPP1-high versus ENPP1-low tumors, we established a clear link between target expression, restored downstream signaling, and tumor efficacy. Specifically, STF-1623 led to tumor cGAMP accumulation, tumor and serum IFN-γ production, and ultimately suppression of tumor only in the EMT6 tumors with high ENPP1 expression and sustained τ but not in MC38 tumors. IFN-γ are pleotropic cytokine with antitumor functions, mainly produced by NK cells, NKT cells, CD8^+^ T cells and CD4^+^ T cells upon activation (*58*). Although STING activation does not directly induce IFN-γ production, it can indirectly promote its expression through downstream immune activation. IFN-γ reflects intratumoral cGAMP-STING activation and can be detected in the serum eight hours within STF-1623 injection. Notably, the serum IFN-γ levels were in the sub-picomolar range, ∼1000 fold below the threshold for IFN-γ receptor binding and activation (*K*_d_ ∼ 0.5nM) (*59*). These results nominate serum INF-γ as an ideal biomarker for predicting STF-1623 treatment responses without causing systemic toxicity.

Prior to this study, the role of ENPP1 in breast cancer has been extensively studied through genetic perturbations (*34,42,43*). Here, we demonstrated the tumor suppression benefits of STF-1623 in combination with existing treatment modalities in pancreatic cancer, colorectal cancer, and glioblastoma, indicating potential utility of STF-1623 across cancer types. Importantly, the versatility of STF-1623 in combination therapies owes to its mechanism of action: it could augment the effect of DNA damaging agents such as alkylating agents and IR by enhancing cGAMP accumulation, as well as synergizing with T cell targeting therapies such as an a-PD-1 and a-PD-L1 by recruiting T cells. Since ENPP1 inhibition work by potentiating the effects of these existing therapies, we reasoned that a short course of STF-1623 would be sufficient to amplify intratumoral cGAMP or recruit TILs, ultimately promoting durable anticancer effects while preventing side-effects. Indeed, most of our studies had a seven-day dosing that was sufficient to reprogram the immune landscape in the tumor microenvironment and elicit tumor suppression while being well-tolerated, demonstrating the superior safety and efficacy of STF-1623.

ENPP1 inhibition has several advantages over the traditional approach of direct STING agonists. The amount of STING agonists delivered intratumorally or systemically is usually in excess. Therefore, exogenous STING agonists floods into and activate both cancer and host cells, resulting in opposing pro-versus anti-cancer effects (*6,16,19–21,34*). Additionally, high levels of STING agonism could result in deleterious effects on tumor vasculatures and T cells (*24–28*), which hamper its therapeutic efficacy. ENPP1 inhibition overcomes these limitations given the target’s distinctive mechanism of activation. First, ENPP1 inhibitor enables the accumulation of endogenous cGAMP produced and released by cancer cells, preventing side-effects due to overstimulation in the case of a bolus injection of STING agonists. Secondly, cancer cells are the cGAMP producing cells (*21,30–32,35*), and most cGAMP transporters shuttles cGAMP down its concentration gradient (*25,36–41*). ENPP1 inhibition restores not only the endogenous levels but the concentration gradient of cGAMP, promoting cGAMP to selectively enter immune cells.

Therefore, our systemically dosed ENPP1 inhibitor can fine tune the level and localization of cGAMP within the tumor microenvironment. It essentially acts as an endogenous tumor TIL-specific STING agonist. Furthermore, ENPP1 promotes cancer through STING-independent pathways, such as by generating the immunosuppressive adenosine (*42,43*) or maintaining cancer stemness (*55*). Therefore. ENPP1 inhibition may provide added benefits than direct STING agonism by interfering with multiple pro-cancer pathways.

Our studies have multiple limitations. We focused on preclinical models and provided limited human data. This is mainly due to technical limitations. Efficacy of ENPP1 inhibition requires a complete immune system that is hard to fully recapitulate with human cancer cell lines or organoids. However, we have good reasons to believe that our preclinical results hold translation values. First, mouse and human ENPP1 are highly conserved in sequence, structure, function, and phenotypes in cancer (*34,52*). Second, STF-1623 inhibits mouse and human ENPP1 with similar potency (*44*) and τ, and we expect it to have similar pharmacokinetic properties in human. Another limitation with our findings is that although we propose that ENPP inhibition is a safer approach to prevent T cell killing from high cGAMP levels (*25–28,34*), our immune characterization in the Panc02 model revealed that STF-1623 administered over 7-day at an intermediate dose (5 mg/kg) induced more CD4^+^ T cell death than 3-day dosing. Nonetheless, 7-day dosing attracted more CD8^+^ T cells, which explain the overall excellent synergy between STF-1623 and a-PD-L1 therapy which mainly depends on CD8^+^ T cells. These results suggest that dosing schedules can be further optimized in clinical trials when combining STF-1623 with checkpoint blockades that depends on CD4^+^ or CD8^+^ T cells respectively.

Our combination of chemical, structural, molecular, and pre-clinical data provide a clear basis for investigating the clinical safety and efficacy of STF-1623. Thoughtful patient selection is key. In the ISPY 2 trail, breast cancer patients whose tumor ENPP1 expression level are within the bottom 50% of the cohort were free of distant metastasis for over 7 years and practically cured after anti-PD-1 neoadjuvant therapy followed by definitive surgery (*34*). This is consistent with the preclinical findings here that tumors with high ENPP1expression are more likely to benefit from the ultralong τ property of STF-1623 and exhibit anti-metastatic efficacy. For example, ENPP1-high EMT6 tumors receiving STF-1623 and anti-PD-L1 combination therapy were free of lung metastasis. For the first STF-1623 clinical trial, we recommend STF-1623 and anti-PD-1/anti-PD-L1 combination therapy in breast patients expressing high ENPP1 levels (activity of which can be measured in the serum) in a neoadjuvant setting. In the trial, we recommend characterizing of serum blood markers that reflect therapeutic response and downstream efficacy such as IFN-γ in our preclinical models to better understand drug action and predict outcomes. In the future, an innate immune checkpoint blocker that achieves tumor specificity like STF-1623 could serve as a potent adjuvant to potentiate various immune therapy modalities including cancer vaccines, immune checkpoint blockade, and CAR-T for solid tumors. With the impressive pre-clinical outcomes, STF-1623 paves ways for the next generation of immunotherapy.

## MATERIALS AND METHODS

### Study design

The study was designed to characterize the pharmacokinetic and pharmacodynamic properties, and preclinical safety and efficacy of an ENPP1 small-molecule inhibitor STF-1623. Systemic and tumor pharmacokinetics of STF-1623 were characterized in various tumor models. Surface plasma resonance was used to characterize dissociation rate and τ of STF-1623 bound to mouse and human ENPP1. A co-crystal structure between STF-1623 and faux human ENPP1 was obtained to examine structural determinants of the compounds potency. A separation-of-function point mutation of ENPP1 revealed that membrane bound ENPP1 in tumors drives tumor progression and constitutes the active target. Intratumoral cGAMP levels upon STF-1623 treatment in mice were measured with mass spectrometry. Safety was assessed by weight loss and change in serum and organ pyrophosphate levels as a marker for on-target side effects. Efficacy of STF-1623 as a single agent and in combination with existing cancer therapies was assessed by primary tumor growth and metastases across breast, colorectal, pancreatic cancers and glioblastoma, supplemented by detailed immune characterization using flow cytometry.

### Surface plasma resonance

N-terminally Avi-tagged and biotinylated mouse or human ENPP1 was immobilized on a SA sensor chip and loaded to about 3000 RU on a Bicaore T200 SPR system. STF-1623 was injected at increasing concentrations (0.12, 0.36, 1.11, 3.33, 10 nM) with a flow rate of 50 µl/min over 200 seconds each in PBS-P+ buffer (Cytiva). One run with mouse ENPP1 used STF-1623 at increasing concentrations (0.062, 0.19, 0.57, 1.67, 5nM) with a flow rate of 30 µl/min. Final dissociation took place over 600 seconds. Target residence time was calculated as the reciprocal of K_off_.

### Crystallization and structure determination

The protein has been purified to homogeneity and concentrated to 10 mg/ml prior to crystallization screening. Fresh protein sample stocks were mixed with inhibitor at 0.12 mM (∼1:1.2 protein:inhibitor molar ratio) prior to mixing with various crystallization buffers from commercial sources. Crystallization experiments were set-up using a Douglas Oryx8 Nanodrop dispensing robot (Douglas Instruments Ltd, Berkshire, United Kingdom). Crystals were grown using the sitting drop vapor diffusion method in an incubator at 16 °C. Crystals were harvested in mother liquor solution supplemented with 25 % glycerol and cryo-cooled by plunging into liquid N_2_. In general, crystals harvested from different crystallization conditions showed a huge variation in X-ray diffracting power and therefore a large number were screened for initial data quality assessment. The best candidates were selected and stored for further data collection. Data collections were performed at cryogenic temperature at Stanford Synchrotron Radiation Lightsource (SSRL) at SLAC National Accelerator Laboratory (Menlo Park, CA, USA) beamlines 9-2 and 12-2 (60). A crystal grown from a 30% PEG 1500 solution was used for full data collection. Data to a Bragg spacing of 2.70 Å were collected at SSRL station BL12-2 using a 15×15 μm microfocused beam. Data was anisotropic and extended to about 2.7-2.8 Å on reciprocal a and c vectors and to about 3.8 Å on reciprocal b vector. The crystal belonged to the tetragonal space group P4_3_2_1_2 and contained one polypeptide chain per asymmetry unit. The structure was solved by the molecular replacement method with Phaser (*61*) using one polypeptide chain of human ENPP3 (PDB ID: 6C02; (*62*)) as the search model stripped from non-protein atoms. The single polypeptide chain encompasses residues Gly52-Phe871 and includes mutated positions Gln244Lys and Glu275Asp. One copy of the inhibitor coordinating one of the zinc ions is in the active site pocket. Cycled with refinement with REFMAC5 (*63*) manual adjustments on the polypeptide chain were made in COOT (*64*). Solvent water molecules, ions, glycans, and the inhibitor molecule were then assigned. Water molecules were placed based on their hydrogen bonding properties. Data was processed with XDS (*65*), DIALS (*66*), AIMLESS (*67*) and analyzed with different computing modules within the CCP4 suite (*68*).Graphic renderings were prepared with pymol. Refinement progressed to convergence and reached an excellent agreement between the model and the experimental data. **Table S3** presents data collection, refinement, and structure quality check parameters.

### STF-1623 efficacy in tumor models

For EMT6 efficacy, EMT6 (5 x 10^5^) were subcutaneously injected into the right flank of 6 weeks old female BALB/c mice. When the average tumor size reached 91 mm^3^, 7 days after tumor inoculation, mice were randomized into 10 per group based on “Matched distribution” method (StudyDirectorTM software, version 3.1.399.19), denoted as day 0 of the study. One day after randomization, the following treatment was administered alone or in combination: STF-1623 (5 mg/kg), subcutaneous, QD day 1-7; anti-PD-L1 (2.5 mg/kg) intraperitoneal, BIW (day 1, 4 each week) x 4 weeks; ionizing radiation (15 Gy), single dose, day 2. Tumors and weight were monitored at least twice weekly. One mouse from STF-1623 + anti-PD-L1 group experience early death on day 8 and was excluded from the study. The study was terminated on day 29, and tumors and lungs were collected for further immunohistochemistry analyses. For EMT6 ionizing radiation dosage optimization, twenty mice with average EMT6 subcutaneous tumor size of ∼75 mm^3^ were randomized 10 days after inoculation, receiving no treatment, or CT-guided focal IR of 5, 10, 15, 20 Gy on the Small Animal Radiation Therapy platform (SmART+). No mortality was observed in this study prior to study termination. For MC38 and CT26 efficacy, MC38 (5 x 10^5^) or CT26 (1 x 10^6^) were subcutaneously injected into the right flank of 6 weeks old female C57BL/6 or BALB/c mice, respectively. When tumor volumes reached ∼90mm^3^ average volume, the mice were stratified into groups of 10 mice, denoted as day 0 of the study. On day 1, the following treatment was administered alone or in combination: STF-1623 (10 mg/kg), subcutaneous, day 1-3, 8-10, 15-17 (MC38 only); anti-PD-1 (3 mg/kg), intraperitoneal, day 1, 4, 8, 11, 15 (MC38 only). Tumors and weight were monitored at least twice weekly. For MC38, one mouse in the vehicle group was found dead on day 17 of the study, and the study was terminated. For CT26, one mouse in the vehicle group had excess tumor burden (>3000 mm3) on day 13, and the study was terminated.

For Panc02 efficacy, Panc02 (3 x 10^6^) were subcutaneously injected into the right flank of 6 weeks old female BALB/c mice. When the average tumor size reached 86 mm^3^, 6 days after tumor inoculation, mice were randomized based on “Matched distribution” method (StudyDirectorTM software, version 3.1.399.19), denoted as day 0 of the study. One day after randomization, the following treatment was administered alone or in combination: STF-1623 (50 or 100 mg/kg) subcutaneous, QD day 1-7; ionizing radiation (20 Gy), single dose, day 2 of the study. Tumors and weight were monitored at least twice weekly. One mouse from ionizing radiation group experienced early death on day 8 and was excluded from the study. The study was terminated no day 49.

For GL261 efficacy: GL261 Red-Fluc (3 x 10^5^) in 3 µL of PBS were intracranially injected into the right frontal lobe (3 mm lateral from the bregma, 0.5 mm from the anterior at a depth of 3.5 mm) of C57BL/6 mice. Four days after tumor inoculation, mice were randomized based on Matched distribution” method (StudyDirectorTM software, version 3.1.399.19), denoted as day 0 of the study. One day after randomization, the following treatment was administered alone or in combination: STF-1623 (50 mg/kg), subcutaneous, QD day 1-7, ionizing radiation (15 Gy), single dose, day 2 of the study, temozolomide (10 mg/kg), per oral, QD, 5 days on, 2 days off x 3 weeks. Tumor growth was checked twice per week by bioluminescent imaging. At 15 minutes prior to imaging, D-Luciferin (PerkinElmer, #122799) was injected intraperitoneally into animal at 150 mg/kg. Anesthetized mice were transferred to the nose cone (ventral) attached to the manifold inside the imaging chamber. Door was closed and the “Acquire” button activated in the living image program (PerkinEler, IVIS Lumina Series III). The order and position (dorsal/ventral, up to 3 mice laid alongside each other in cage order) remained constant throughout study. Duration and binning (sensitivity) of the image was dependent upon the intensity of the lesions present. All study groups without temozolomide were terminated on study day 21 for ethical considerations; all study groups with temozolomide were terminated on study day 46.

### Liquid chromatography and mass spectrometry (LC-MS)

For detection of STF-1623 (**Fig. 1B,C**) and cGAMP (**Fig. 4A, S4A**) in EMT6 and MC38 studies, EMT6 or MC38 (1 x 10^6^) were subcutaneously injected into female BALB/c or C57BLJ/6 mice, respectively. Upon tumor establishment (70-95 mm^3^), one dose of STF-1623 (50 mg/mL) or vehicle was injected subcutaneously. Whole tumor and terminal serum samples were collected at 0.25, 0.5, 2, 4, 6, 8, 24, 48 hours (n = 3 mice/time point). For STF-1623 measurement, 2 µL of prepared samples were analyzed on an API-4000Qtrap mass spectrometer, ESI positive, MRM scan with a Shimadzu HPLC/Autosampler with ACE C8 column (2.1 x 50mm, 5 µm). For cGAMP measurement, 15 µL of prepared samples analyzed on an API-4000Qtrap mass spectrometer, ESI positive, MRM scan with a Shimadzu HPLC/Autosampler with Phenomenex Luna C18 column (100 x 2 mm). The mobile phase consisted of 5 mM ammonium acetate and 1% formic acid (A) and acetonitrile (100%) and 1% formic acid (B).

For **Fig. 6A**, STF-1623 (50 or 100 mg/kg) was administered subcutaneously to female C57BL/6 mice. Terminal blood sampling was performed at 0.083, 0.25, 1, 2, 4, 8, and 24 hours (n = 3 mice/time point). Whole brain and terminal cerebrospinal fluid (CSF) samples were collected from each mouse at termination. Mice were perfused with cold PBS prior to harvesting brains tissues. 2 µL of prepared samples were analyzed on an API-4000Qtrap mass spectrometer, ESI positive, MRM scan with a Shimadzu HPLC/Autosampler with ACE C8 column (2.1×50mm, 5 µm). The mobile phase consisted of 5 mM ammonium acetate and 1% formic acid (A) and acetonitrile (100%) and 1% formic acid (B).

For detection of STF-1623 in 4T1 tumor and serum (**Fig S1A**), WT or *Enpp1^-/-^* 4T1 (5 x 10^4^) were orthotopically injected into female BALB/c mice. Upon tumor establishment, one dose of STF-1623 (50 mg/mL) or vehicle was injected subcutaneously. Whole tumor and terminal serum samples were collected at 0.5, 2, 4, 8, 24 hours (n = 2-3 mice/time point). Samples are analyzed following steps described in a previous paper (44). Briefly, organ homogenates and serum were precipitated with acetonitrile, centrifuged at 16,000 x *g*, and resuspended in a matrix of 2:1 0.1% formic acid:acetonitrile with clemizole as the internal standard. LC-MS was performed on a Shimadzu HPLC with an autosampler set at 4 °C and connected to an AB Sciex 4000.

For detection of STF-1623 in healthy tissues (**Fig. S1B**), one dose of STF-1623 (50 mg/kg) was subcutaneously injected into WT or *Enpp1^-/-^*BALB/c mice. Liver, kidney and terminal serum samples were collected at 0.5, 2, 4, 8, 24 hours (n = 2-3 mice/time point). 20 µL serum samples were directly loaded to a 96-well Millipore Multiscreen Solvinert 0.45 micron low binding PTFE hydrophilic filter plate. Tissue samples were homogenized with water (x3 dilution) then 20 µL was loaded to the filter plate. All plasma/tissue samples were treated with 60 µL 90/10 acetonitrile/water with Carbamazepine as internal standard to extract the analyte and precipitate protein. The plates were agitated on ice for approximately ten minutes prior to centrifugation into a collection plate. Separate standard curves were prepared in blank mouse plasma and tissue homogenate and processed in parallel with the samples. The filtrate was directly analyzed by LC-MS/MS analysis against. HPLC and MS/MS parameters are provided in **Table S4**.

### Quantification and Statistical Analysis

In ENPP1/3 inhibition assays, the half maximal inhibitory concentrations (IC_50_) were obtained by sigmoidal dose-response fitting with Prism software. In cGAMP degradation assays and pharmacokinetics analyses, half-life was obtained by one phase exponential decay fitting with Prism software. Enzyme kinetics data are fit with Michaelis-Menten model with Prism software. Graphs show means and standard deviation (± SD) or standard error of the mean (± SEM). Statistical significance, group size, and experimental details are described in the figure legends.

## List of Supplementary Materials

Present a list of the Supplementary Materials in the following format.

## Materials and Methods

Fig S1 to S6

Tables S1 to S4

Source Data Fig 1 to 3

References (*69–71*)

## Acknowledgments

We thank B. Plosky and the rest of the Scientific Publication Team at Arc Institute for constructive feedback on the manuscript. We thank all other Li Lab members for their constructive comments and discussion through the course of this study. Use of the Stanford Synchrotron Radiation Lightsource, SLAC National Accelerator Laboratory, is supported by the U.S. Department of Energy, Office of Science, Office of Basic Energy Sciences under Contract No. DE-AC02-76SF00515. The SSRL Structural Molecular Biology Program is supported by the DOE Office of Biological and Environmental Research, and by the National Institutes of Health, National Institute of General Medical Sciences (P30GM133894). The contents of this publication are solely the responsibility of the authors and do not necessarily represent the official views of NIGMS or NIH.

## Funding

National Institute of Health grant 5R01CA258427-02 (LL)

Arc Institute (LL, SW, JAC, DF, REM)

Angarus Therapeutics (RMJ, NV, JS, JP, NR)

National Institute of Health grant P30GM133894 (DF)

## Author contributions

Conceptualization: RMJ, NV, JS, JP, NR, LL

Methodology: RMJ, SW, JAC, DF, REM, NV, JS, JP,

Investigation: SW, JAC, DF, REM, XC

Visualization: SW, JAC, LL

Funding acquisition: NR, LL

Supervision: RMJ, LL

Writing – original draft: RMJ, SW, JAC, DF, LL

Writing – review & editing: SW, LL

## Data and materials availability

Human faux ENPP1 with STF-1623 structure was deposited (PDB: 9NIR). All unique reagents generated in this study are available from the lead contact with a completed Materials Transfer Agreement. All data reported in this paper will be shared by the lead contact upon request. Any additional information required to reanalyze the data reported in this paper is available from the lead contact upon request. Further information and requests for resources and reagents should be directed to and will be fulfilled by the lead contact, Lingyin Li.

## References

1. Miller KD, Nogueira L, Devasia T, Mariotto AB, Yabroff KR, Jemal A, et al. Cancer treatment and survivorship statistics. CA Cancer J Clin. 2022;72(5):409–36.

2. Haslam A, Olivier T, Prasad V. How many people in the US are eligible for and respond to checkpoint inhibitors: An empirical analysis. Int J cancer. 2025.

3. Leon-Ferre RA, Jonas SF, Salgado R, Loi S, De Jong V, Carter JM, et al. Tumor-Infiltrating Lymphocytes in Triple-Negative Breast Cancer. JAMA. 2024;331(13):1135– 44.

4. Bonaventura P, Shekarian T, Alcazer V, Valladeau-Guilemond J, Valsesia-Wittmann S, Amigorena S, et al. Cold Tumors: A Therapeutic Challenge for Immunotherapy. Front Immunol. 2019.

5. Wu B, Zhang B, Li B, Wu H, Jiang M. Cold and hot tumors: from molecular mechanisms to targeted therapy. Signal Transduct Target Ther. 2024;9(1):274.

6. Deng L, Liang H, Xu M, Yang X, Burnette B, Arina A, et al. STING-dependent cytosolic DNA sensing promotes radiation-induced type I interferon-dependent antitumor immunity in immunogenic tumors. Immunity. 2014;41(5):843–52.

7. Yum S, Li M, Fang Y, Chen ZJ. TBK1 recruitment to STING activates both IRF3 and NF-κB that mediate immune defense against tumors and viral infections. Proc Natl Acad Sci U S A. 2021;118(14):e2100225118.

8. Wu J, Dobbs N, Yang K, Yan N. Interferon-Independent Activities of Mammalian STING Mediate Antiviral Response and Tumor Immune Evasion. Immunity. 2020 Jul 14;53(1):115–126.e5.

9. Ishikawa H, Barber GN. STING is an endoplasmic reticulum adaptor that facilitates innate immune signalling. Nat. 2008;455(7213):674–8.

10. Ablasser A, Goldeck M, Cavlar T, Deimling T, Witte G, Röhl I, et al. cGAS produces a 2′-5′-linked cyclic dinucleotide second messenger that activates STING. Nat. 2013;498(7454):380–4.

11. Gao P, Ascano M, Wu Y, Barchet W, Gaffney BL, Zillinger T, et al. Cyclic [G(2′,5′)pA(3′,5′)p] Is the Metazoan Second Messenger Produced by DNA-Activated Cyclic GMP-AMP Synthase. Cell. 2013 May 23;153(5):1094–107.

12. Wu J, Sun L, Chen X, Du F, Shi H, Chen C, et al. Cyclic GMP-AMP is an endogenous second messenger in innate immune signaling by cytosolic DNA. Science. 2013;339(6121):826–30.

13. Sun L, Wu J, Du F, Chen X, Chen ZJ. Cyclic GMP-AMP synthase is a cytosolic DNA sensor that activates the type I interferon pathway. Science. 2013;339(6121):786–91.

14. Chan R, Cao X, Ergun SL, Njomen E, Lynch SR, Ritchie C, et al. Human STING oligomer function is governed by palmitoylation of an evolutionarily conserved cysteine. bioRxiv. 2024;2023.08.11.553045.

15. Ergun SL, Fernandez D, Weiss TM, Li L. STING Polymer Structure Reveals Mechanisms for Activation, Hyperactivation, and Inhibition. Cell. 2019;178(2):290–301.e10.

16. Woo SR, Fuertes MB, Corrales L, Spranger S, Furdyna MJ, Leung MYK, et al. STING-dependent cytosolic DNA sensing mediates innate immune recognition of immunogenic tumors. Immunity. 2014;41(5):830–42.

17. Meric-Bernstam F, Sweis RF, Kasper S, Hamid O, Bhatia S, Dummer R, et al. Combination of the STING Agonist MIW815 (ADU-S100) and PD-1 Inhibitor Spartalizumab in Advanced/Metastatic Solid Tumors or Lymphomas: An Open-Label, Multicenter, Phase Ib Study. Clin Cancer Res. 2023;29(1):110–21.

18. Meric-Bernstam F, Sweis RF, Hodi FS, Messersmith WA, Andtbacka RHI, Ingham M, et al. Phase I Dose-Escalation Trial of MIW815 (ADU-S100), an Intratumoral STING Agonist, in Patients with Advanced/Metastatic Solid Tumors or Lymphomas. Clin Cancer Res. 2022;28(4):677–88.

19. Wang Q, Bergholz JS, Ding L, Lin Z, Kabraji SK, Hughes ME, et al. STING agonism reprograms tumor-associated macrophages and overcomes resistance to PARP inhibition in BRCA1-deficient models of breast cancer. Nat Commun. 2022;13(1):1–17.

20. Li J, Hubisz MJ, Earlie EM, Duran MA, Hong C, Varela AA, et al. Non-cell-autonomous cancer progression from chromosomal instability. Nat. 2023;620(7976):1080–8.

21. Bakhoum SF, Ngo B, Laughney AM, Cavallo JA, Murphy CJ, Ly P, et al. Chromosomal instability drives metastasis through a cytosolic DNA response. Nature. 2018;553(7689):467–72.

22. Yang H, Lee WS, Kong SJ, Kim CG, Kim JH, Chang SK, et al. STING activation reprograms tumor vasculatures and synergizes with VEGFR2 blockade. J Clin Invest. 2019;129(10):4350.

23. Wang L, Liang Z, Guo Y, Habimana J de D, Ren Y, Amissah OB, et al. STING agonist diABZI enhances the cytotoxicity of T cell towards cancer cells. Cell Death Dis. 2024;15(4):1–11.

24. Downey CM, Aghaei M, Schwendener RA, Jirik FR. DMXAA Causes Tumor Site-Specific Vascular Disruption in Murine Non-Small Cell Lung Cancer, and like the Endogenous Non-Canonical Cyclic Dinucleotide STING Agonist, 2′3′-cGAMP, Induces M2 Macrophage Repolarization. PLoS One. 2014;9(6):e99988.

25. Sudaryo V, Carvalho DR, Lee JM, Carozza JA, Cao X, Cordova AF, et al. Toxicity of extracellular cGAMP and its analogs to T cells is due to SLC7A1-mediated import. bioRxiv. 2024;2024.09.21.614248.

26. Sivick KE, Desbien AL, Glickman LH, Reiner GL, Corrales L, Surh NH, et al. Magnitude of Therapeutic STING Activation Determines CD8+ T Cell-Mediated Anti-tumor Immunity. Cell Rep. 2018;25(11):3074–3085.e5.

27. Gulen MF, Koch U, Haag SM, Schuler F, Apetoh L, Villunger A, et al. Signalling strength determines proapoptotic functions of STING. Nat Commun. 2017;8(1):1–10.

28. Larkin B, Ilyukha V, Sorokin M, Buzdin A, Vannier E, Poltorak A. Cutting Edge: Activation of STING in T Cells Induces Type I IFN Responses and Cell Death. J Immunol. 2017;199(2):397–402.

29. Li L, Yin Q, Kuss P, Maliga Z, Millán JL, Wu H, et al. Hydrolysis of 2′3′-cGAMP by ENPP1 and design of nonhydrolyzable analogs. Nat Chem Biol. 2014;10(12):1043–8.

30. MacKenzie KJ, Carroll P, Martin CA, Murina O, Fluteau A, Simpson DJ, et al. cGAS surveillance of micronuclei links genome instability to innate immunity. Nat. 2017;548(7668):461–5.

31. Flynn PJ, Koch PD, Mitchison TJ. Chromatin bridges, not micronuclei, activate cGAS after drug-induced mitotic errors in human cells. Proc Natl Acad Sci U S A. 2021;118(48).

32. Chen YA, Shen YL, Hsia HY, Tiang YP, Sung TL, Chen LY. Extrachromosomal telomere repeat DNA is linked to ALT development via cGAS-STING DNA sensing pathway. Nat Struct Mol Biol. 2017;24(12):1124–31.

33. Konno H, Yamauchi S, Berglund A, Putney RM, Mulé JJ, Barber GN. Suppression of STING signaling through epigenetic silencing and missense mutation impedes DNA damage mediated cytokine production. Oncogene. 2018;37(15):2037–51.

34. Wang S, Böhnert V, Joseph AJ, Sudaryo V, Skariah G, Swinderman JT, et al. ENPP1 is an innate immune checkpoint of the anticancer cGAMP–STING pathway in breast cancer. Proc Natl Acad Sci U S A. 2023;120(52):e2313693120.

35. Carozza JA, Böhnert V, Nguyen KC, Skariah G, Shaw KE, Brown JA, et al. Extracellular cGAMP is a cancer-cell-produced immunotransmitter involved in radiation-induced anticancer immunity. Nat Cancer. 2020;1(2):184–96.

36. Ritchie C, Cordova AF, Hess GT, Bassik MC, Li L. SLC19A1 Is an Importer of the Immunotransmitter cGAMP. Mol Cell. 2019;75(2):372–381.e5.

37. Luteijn RD, Zaver SA, Gowen BG, Wyman SK, Garelis NE, Onia L, et al. SLC19A1 transports immunoreactive cyclic dinucleotides. Nature. 2019;573(7774):434–8.

38. Cordova AF, Ritchie C, Böhnert V, Li L. Human SLC46A2 Is the Dominant cGAMP Importer in Extracellular cGAMP-Sensing Macrophages and Monocytes. ACS Cent Sci. 2021;7(6):1073–88.

39. Lahey LJ, Mardjuki RE, Wen X, Hess GT, Ritchie C, Carozza JA, et al. LRRC8A:C/E Heteromeric Channels Are Ubiquitous Transporters of cGAMP. Mol Cell. 2020;80(4):578–591.e5.

40. Concepcion AR, Wagner LE, Zhu J, Tao AY, Yang J, Khodadadi-Jamayran A, et al. The volume-regulated anion channel LRRC8C suppresses T cell function by regulating cyclic dinucleotide transport and STING-p53 signaling. Nat Immunol. 2022;23(2):287–302.

41. Zhou Y, Fei M, Zhang G, Liang WC, Lin WY, Wu Y, et al. Blockade of the Phagocytic Receptor MerTK on Tumor-Associated Macrophages Enhances P2X7R-Dependent STING Activation by Tumor-Derived cGAMP. Immunity. 2020;52(2):357–373.e9.

42. De Córdoba BRF, Moreno H, Valencia K, Perurena N, Ruedas P, Walle T, et al. Tumor ENPP1 (CD203a)/Haptoglobin Axis Exploits Myeloid-Derived Suppressor Cells to Promote Post-Radiotherapy Local Recurrence in Breast Cancer. Cancer Discov. 2022;12(5):1356.

43. Li J, Duran MA, Dhanota N, Chatila WK, Bettigole SE, Kwon J, et al. Metastasis and Immune Evasion from Extracellular cGAMP Hydrolysis. Cancer Discov. 2021;11(5):1212–27.

44. Carozza JA, Brown JA, Böhnert V, Fernandez D, AlSaif Y, Mardjuki RE, et al. Structure-Aided Development of Small-Molecule Inhibitors of ENPP1, the Extracellular Phosphodiesterase of the Immunotransmitter cGAMP. Cell Chem Biol. 2020;27(11):1347–1358.e5.

45. Zeng Z, Wong CJ, Yang L, Ouardaoui N, Li D, Zhang W, et al. TISMO: syngeneic mouse tumor database to model tumor immunity and immunotherapy response. Nucleic Acids Res. 2022;50(D1):D1391–7.

46. Stabach PR, Zimmerman K, Adame A, Kavanagh D, Saeui CT, Agatemor C, et al. Improving the Pharmacodynamics and In Vivo Activity of ENPP1-Fc Through Protein and Glycosylation Engineering. Clin Transl Sci. 2021;14(1):362–72.

47. Davis MJ, Hanson KA, Clark F, Fink JL, Zhang F, Kasukawa T, et al. Differential use of signal peptides and membrane domains is a common occurrence in the protein output of transcriptional units. PLoS Genet. 2006;2(4):554–63.

48. Jansen S, Perrakis A, Ulens C, Winkler C, Andries M, Joosten RP, et al. Structure of NPP1, an Ectonucleotide Pyrophosphatase/Phosphodiesterase Involved in Tissue Calcification. Structure. 2012 Nov 7;20(11):1948–59.

49. Teufel F, Almagro Armenteros JJ, Johansen AR, Gíslason MH, Pihl SI, Tsirigos KD, et al. SignalP 6.0 predicts all five types of signal peptides using protein language models. Nat Biotechnol. 2022;40(7):1023–5.

50. Sakaguchi M, Tomiyoshi R, Kuroiwa T, Mihara K, Omura T. Functions of signal and signal-anchor sequences are determined by the balance between the hydrophobic segment and the N-terminal charge. Proc Natl Acad Sci U S A. 1992;89(1):16.

51. Fabian KP, Padget MR, Fujii R, Schlom J, Hodge JW. Differential combination immunotherapy requirements for inflamed (warm) tumors versus T cell excluded (cool) tumors: engage, expand, enable, and evolve. J Immunother cancer. 2021;9(2).

52. Carozza JA, Cordova AF, Brown JA, AlSaif Y, Bohnert V, Cao X, et al. ENPP1’s regulation of extracellular cGAMP is a ubiquitous mechanism of attenuating STING signaling. Proc Natl Acad Sci U S A. 2022;119(21).

53. Orriss IR, Arnett TR, Russell RGG. Pyrophosphate: a key inhibitor of mineralisation. Curr Opin Pharmacol. 2016;28:57–68.

54. Mardjuki R, Wang S, Carozza J, Zirak B, Subramanyam V, Abhiraman G, et al. Identification of the extracellular membrane protein ENPP3 as a major cGAMP hydrolase and innate immune checkpoint. Cell Rep. 2024;43(5).

55. Bageritz J, Puccio L, Piro RM, Hovestadt V, Phillips E, Pankert T, et al. Stem cell characteristics in glioblastoma are maintained by the ecto-nucleotidase E-NPP1. Cell Death Differ. 2014;21(6):929–40.

56. Rudalska R, Harbig J, Forster M, Woelffing P, Esposito A, Kudolo M, et al. First-in-class ultralong-target-residence-time p38α inhibitors as a mitosis-targeted therapy for colorectal cancer. Nat Cancer. 2025;6.

57. De Cesco S, Kurian J, Dufresne C, Mittermaier AK, Moitessier N. Covalent inhibitors design and discovery. Eur J Med Chem. 2017;138:96–114.

58. Jorgovanovic D, Song M, Wang L, Zhang Y. Roles of IFN-γin tumor progression and regression: A review. Biomark Res. 2020;8(1):1–16.

59. Kemna J, Gout E, Daniau L, Lao J, Weißert K, Ammann S, et al. IFNγ binding to extracellular matrix prevents fatal systemic toxicity. Nat Immunol 2023 243. 2023;24(3):414–22.

60. Russi S, Song J, Mcphillips SE, Cohen AE. The Stanford Automated Mounter: pushing the limits of sample exchange at the SSRL macromolecular crystallography beamlines. J Appl Crystallogr. 2016;49(Pt 2):622–6.

61. McCoy AJ, Grosse-Kunstleve RW, Adams PD, Winn MD, Storoni LC, Read RJ. Phaser crystallographic software. J Appl Crystallogr. 2007;40(Pt 4):658–74.

62. Gorelik A, Randriamihaja A, Illes K, Nagar B. Structural basis for nucleotide recognition by the ectoenzyme CD203c. FEBS J. 2018 Jul 1;285(13):2481–94.

63. Murshudov GN, Vagin AA, Dodson EJ. Refinement of macromolecular structures by the maximum-likelihood method. Acta Crystallogr D Biol Crystallogr. 1997;53(Pt 3):240–55.

64. Emsley P, Lohkamp B, Scott WG, Cowtan K. Features and development of Coot. Acta Crystallogr D Biol Crystallogr. 2010;66(Pt 4):486–501.

65. Kabsch W. XDS. Acta Crystallogr D Biol Crystallogr. 2010;66(Pt 2):125–32.

66. Winter G, Waterman DG, Parkhurst JM, Brewster AS, Gildea RJ, Gerstel M, et al. DIALS: implementation and evaluation of a new integration package. Acta Crystallogr Sect D, Struct Biol. 2018;74(Pt 2):85–97.

67. Evans PR. An introduction to data reduction: space-group determination, scaling and intensity statistics. Acta Crystallogr Sect D Biol Crystallogr. 2011;67(Pt 4):282.

68. Winn MD, Ballard CC, Cowtan KD, Dodson EJ, Emsley P, Evans PR, et al. Overview of the CCP4 suite and current developments. Acta Crystallogr D Biol Crystallogr. 2011;67(Pt 4):235–42.

69. Kato K, Nishimasu H, Okudaira S, Mihara E, Ishitani R, Takagi J, et al. Crystal structure of Enpp1, an extracellular glycoprotein involved in bone mineralization and insulin signaling. Proc Natl Acad Sci U S A . 2012 Oct 16;109(42):16876–81.

70. Mardjuki RE, Carozza JA, Li L. Development of cGAMP-Luc, a sensitive and precise coupled enzyme assay to measure cGAMP in complex biological samples. J Biol Chem. 2020;295(15):4881–92.

71. Dennis ML, Newman J, Dolezal O, Hattarki M, Surjadi RN, Nuttall SD, et al. Crystal structures of human ENPP1 in apo and bound forms. Acta Crystallogr Sect D Struct Biol. 2020;76(9):889–98.

