## Supplemental Information for "ENPP1 inhibitor with ultralong drug-target residence time as an innate immune checkpoint blockade cancer therapy"

**Materials and Methods**

**Mouse strains**

C57BL/6 and BALB/c mice were purchased from the Jackson Laboratory, Charles River, or Taconic Biosciences. BALB/cJ-*Enpp1^asj-2J/GrsrJ^* (*Enpp1^-/-^,* strain #019107) was purchased from the Jackson Laboratory. For all tumor experiments, female mice around 6 weeks old were used. Mice maintained at Stanford University followed the Stanford University Institutional Animal Care and Use Committee (IACUC) regulations. All procedures done at Stanford were approved by the Stanford University Administrative Panel on Laboratory Animal Care (APLAC).

**Mammalian cell lines**

293T *ENPP1*^-/-^ cells were generated in a previous study (*35*). Expi293F cells were procured from ThermoFisher. 4T1, EMT6, Panc02, CT26 cells were procured from ATCC. MC38 cells were procured from Kerfast. IVISbrite GL261 Red-Fluc (GL261) cells were procured from Revvity. 4T1 *Enpp1*^-/-^, 4T1 ENPP1^WT-OE^, and ENPP^T238A-OE^ cells were generated in a previous study (*34*). 4T1, MC38. CT26, Panc02, and their derived cell lines were maintained in RPMI (Corning Cellgro) supplemented with 10% FBS (R&D Systems), 10 mM HEPES (Gibco), and 1% penicillin-streptomycin (ThermoFisher). 293T *ENPP1^-/-^*, EMT6, GL261 cells were maintained in DMEM (Corning Cellgro) supplemented with 10% FBS and 1% penicillin-streptomycin. Expi293F cells (Thermo Fisher) were were maintained in a 2:1 mixture of FreeStyle 293 Medium:Expi293 Expression Medium at 37°C, 80% humidity, 8% CO_2_, shaking at 120 rpm. All cells were maintained in a humidified incubator at 37°C and 5% CO_2_. All cell lines tested negative for mycoplasma contamination.

**Synthesis of STF-1623 (2-(1-(8-methoxyquinazolin-4-yl)piperidin-4-yl)ethyl)phosphonic acid)**

STF-1623-acid (denoted as STF-1623 throughout) was synthesized using a modified procedure from steps described in a previous study (*44*). Briefly, 1,8-Diazabicyclo[5.4.0]undec-7-ene (DBU) was added to a solution of dibenzyl 2‐(piperidin‐4‐yl)ethylphosphonate TFA salt (1.0 mole eq.) in 1,4 dioxane, held at NMT 5°C and over NLT 30 min. 4‐chloro‐8‐methoxyquinazoline (1.0 mole eq.) was then added over a time period so that process temperature was below 10°C and stirred overnight until reaction completion. The reaction mixture was worked up by dilution with water and extracted with 2-methyl tetrahydrofuran (2‑Me-THF). As part of the work up and isolation procedure, dibenzylphosphonate (Compound 6) was converted to the methanesulfonic acid salt *in situ* by diluting the combined organic phases with water and the addition of methane sulfonic acid until the pH was not more than 1.0. The aqueous phase was washed with 2-Me-THF then freebased by the addition of sodium hydroxide pellets until the pH was 4.0 ± 0.2. Freebased Compound 6 was extracted with 2-Me-THF, azeotroped 3 times with 2‑Me‑THF, then evaporated to yield Compound 6 as an oil. The potency of Compound 6 was determined by qNMR. A solution of the Compound 6 in methanol was flushed with nitrogen then treated with 10% Palladium on Carbon, the reactor purged with hydrogen then pressurized to 20 inches of water of hydrogen pressure and agitated to create a large vortex. Completion of reaction was determined by HPLC (~3 hours). The reaction mixture was filtered through two Meissner cartridge filters set up in series (30 μm then 0.45 μm) and the solvent removed by distillation. Water was added through a 30 μm Meissner filter and distillation continued until the volume of the mixture has been reduced to a constant volume. The resulting mixture was slurried with water at 60°C for 3 h then the cooling ramped to 20 °C over 4 h and held at 20 °C for 12 h. The slurry was filtered, washed with water and dried under a stream of nitrogen, then under full vacuum at 50 °C for 12 hours to give STF-1623-acid as a dihydrate. STF-1623 was stored at room temperature.

**Recombinant DNA**

Following the same strategy as previously published secreted ENPP expression constructs (*69*), secreted human ENPPs were created by fusing the N-terminal secretion sequence/furin cleavage sequence of mouse ENPP2 (residues 1-58) with human ENPP1 (residues 110-925) or human ENPP3 (residues 55-875). These ENPP sequences were inserted into a pcDNA3 plasmid containing a C-terminal His12 tag to generate pcDNA3-sec-hENPP1-His12 and pcDNA3-sec-hENPP3-His12. To clone pcDNA3-sec-hENPP3-Q244K/E275D-His12 (faux ENPP1), primer sequences carrying the desired point mutations (**Table S2**) were introduced into pcDNA3-sec-hENPP3-His12 plasmid using QuickChange mutagenesis. To clone pcDNA3-mENPP1-mutant-Flag, primer sequences carrying the desired point mutations (**Table S2**) were introduced into pcDNA3-mENPP1-WT-Flag (Genescript) using QuickChange mutagenesis. To clone pLenti-CMV-mENPP1-A84S-Puro, mENPP1-WT sequence was amplified from pcDNA3-mENPP1A84S-Flag using pLenti_mENPP1_fwd and pLenti_mENPP1_rev primers in **Table S2** and inserted into the XbaI-BamHI sites of pLenti-CMV-GFP-Puro (Addgene).

**Generation of stable expression cell lines**

To generate ENPP1^A84S-OE^ cell line, 4T1 *Enpp1^-/-^* cells were virally transduced to stably express A84S mouse ENPP1. Briefly, lentiviral packaging plasmids (pHDM-G, pHDM-Hgmp2, pHDM-tat1b, and pRC.CMV-rev1b) were purchased from Harvard Medical School. 500 ng of pLenti-CMV-mENPP1-A84S-GFP-Puro and 500 ng of each of the packaging plasmids were transfected into 293T cells using FuGENE 6 transfection reagent (Promega). The viral media was exchanged after 24 h, harvested after 48 h and passed through a 0.45 μm filter, and used to transduce 4T1 *Enpp1^-/-^* cells. 48 hours later, cells were selected with 1–2 μg/ml puromycin, single-cell cloned, and 4-6 clones were pooled after verification by western blot and activity assay.

**Preparation of cell lysate and supernatant**

293T *ENPP1^-/-^* cells were transiently transfected with 1 μg WT or mutant, mouse or human pcDNA-ENPP1-FLAG or pLenti-CMV-hENPP3-WT-FLAG plasmids in 3 μL FuGENE 6 (Progrema) in 200 μL Opti-MEM (Thermo Fischer Scientific). One day later, cells were replaced with serum free DMEM media, as ENPP1 in the FBS would interfere with activity read out. The next day, supernatant was collected, span down at 1000 g for 5 minutes to get rid of dead cells and debris, and filtered through an Amicon 30K filter (Sigma-Aldrich) at 12,000 g for 20-30 minutes at 4 °C, until the concentrate volume equals to that of the lysate so ENPP1 expression and activity from the two fractions can be compared directly. Cells are lysed in 50-100 μL of lysis buffer (10 mM Tris pH 9.0, 150 mM NaCl, 10 μM ZnCl_2_, 1% NP-40). For activity assay, supernatant and lysate samples are used as is. For western blotting, 30 μL samples are mixed with 5x reducing or non-reducing sample buffer, heated at 95 °C or five minutes, and sonicated.

**Expression and purification of ENPP proteins**

hENPP1, hENPP3, and faux hENPP1 (hENPP3-Q244D/E275D) in **Fig. 2** were expressed inExpi293F cells. One day prior to transfection, the cells were split to 3 x 10^6^ cells/mL in baffled flasks (Corning). On the day of transfection, cells were diluted to 3-4 x 10^6^ cells/mL (if not already within the range) and transfected with plasmid DNA (0.5 μg DNA/mL cells) using FectoPro (Polyplus) (1 μL FectoPro/mL cells). Cells were immediately boosted with valproic acid (3 μM) and D-glucose (4 g/L). Cells were cultured for an additional 3-5 days. The media was harvested by centrifuging at 3000 x *g* for 10 minutes and passing through a 0.45 μm filter. Media was diluted 1:1 with PBS and imidazole was added to a final concentration of 5 mM. This solution was batch bound with HisPur cobalt resin (Thermo Fisher) (1 mL resin/ 60 mL culture) for 1 hour at 4 °C then loaded onto a fritted column. The column was washed two times with 10 column volumes (CV) wash buffer (PBS + 10 mM imidazole). Eight elutions were performed each with 1 CV elution buffer (PBS + 250 mM imidazole). Elution fractions containing protein were pooled and dialyzed against dialysis buffer (20 mM Tris pH 7.4, 150 mM NaCl, 0.5 mM CaCl_2_, 10 μM ZnCl_2_) overnight at 4 °C. Protein was concentrated to >1 mg/mL, then diluted into 1 μM aliquots supplemented with 0.1% NP-40, and snap frozen for storage at -80 °C.

Transmembrane and secreted mENPP1 in **Fig. 3** are purified as following. 293T *ENPP1^-/-^* cells in a 70% confluent 10 cm^2^ dish were transiently transfected with 5 μg pcDNA-mENPP1-WT-FLAG plasmid in 15 μL FuGENE 6 (Progrema) in 1 mL Opti-MEM (Thermo Fischer Scientific). The next day, cells were passaged to 15 cm^2^ dish in serum free media. The next day, supernatant was collected, span down at 1,000 x *g* for 5 minutes to get rid of dead cells and debris, and concentrated through an Amicon 30K filter (Sigma-Aldrich) at 3,000 x *g* for 15 minutes at 4 °C. Cells were scraped off the plate in 10 mL PBS, spun down at 1,000 x *g* for 10 minutes, and cell pellets were snap frozen in liquid nitrogen. Cell pellet was thawed in 1 mL hypotonic buffer (10 mM HEPES pH 7.5, 25 mM NaCl) on ice for 20 minutes, homogenized with a dounce for 30 strokes, and span down at 23,000 x *g* for 15 minutes at 4 °C. Pellet was solubilized with solubilization buffer (25 mM HEPES pH 7.5, 300 mM NaCl, 1% DDM0.1% CHS) for 1 hour at 4 °C. Upon spinning down at 23,000 g for 30 minutes, supernatant was collected. Both media and lysate fraction then underwent purification with anti-DYKDDDDK (FLAG) magnetic agarose beads (ThermoFischer) overnight at 4 °C, and eluted with 200 μg/mL 3xFLAG peptide (Sigma) in 50 μL elution buffer (25 mM HEPES pH7.5, 150 mM NaCl, 0.1% DDM/0.01% CHS) for five times. Pooled elutions were buffer exchanged with column. Protein concentration was measured with Pierce^TM^ 660nm protein assay reagent (ThermoFisher) and diluted to 40 nM.

**[^32^P] cGAMP activity assay by thin-layer chromatography assay**

cGAMP activity assay (total volume 10-20 μl) containing 50% cell lysate or supernatant, cGAMP (1 μM, with trace [^32^P] cGAMP spiked in), and ENPP1 activity buffer (50 mM Tris pH 9, 250 mM NaCl, 0.5 mM CaCl_2_, 1 μM ZnCl_2_) took place in room temperature cGAMP activity assay (total volume 10-20 μL) containing 50% tumor lysate or serum from mice, cGAMP (1 μM, with trace [^32^P] cGAMP spiked in) took place at 37 °C. To characterize the kinetics of transmembrane or secreted mouse ENPP1, 10 nM of either enzyme (purification following steps described in the section immediate above) was incubated with the indicated concentrations of cGAMP spiked with [^32^P] cGAMP in ENPP1 activity buffer at room temperature. At indicated times, 1 μl aliquots of the reaction were quenched by spotting on HP-TLC silica gel plates (Millipore, Cat# 1.05548.0001). The TLC plates were run in mobile phase (85% ethanol, 5 mM NH_4_HCO_3_) and exposed to a phosphor screen (GE BAS-IP MS). Screens were imaged on a Typhoon 9400 scanner and the ^32^P signal was quantified using ImageJ.

**cGAMP activity assay by cGAMP-Luc**

ENPP1 (15 pM) or ENPP3 and fxENPP1 (250 pM) were added to cGAMP (2 μM). Reactions were incubated and then heat inactivated at 95 °C for 10 minutes. The AMP degradation product was converted to ATP using an enzyme mixture of polyphosphate:AMP phosphotransferase (PAP) and myokinase (MilliporeSigma), which was detected using luciferase (CellTiterGlo, Promega) according to previous publication (*70*). Specific steps are describe in a previous publication (*44*).

**ATP activity assay by CellTiterGlo**

ATP activity assays (10 μL total in a 384 well PCR plate) were composed of the following: cell lysate/supernatant (1%), 1 μM ATP (Sigma) and ENPP1 activity buffer (50 mM Tris pH 9, 250 mM NaCl, 0.5 mM CaCl_2_, 1 μM ZnCl_2_). Reactions were started at indicated times and ended simultaneously by heating at 95 °C for 10 minutes. Reactions (5 μL) were transferred to a white 384 well plate, mixed with CellTiterGlo (5 μL), and luminescence was read after 15 minutes on a Tecan Spark plate reader.

**Western blotting**

Cell lysates or supernatants in reducing or non-reducing sample buffer were separated on an SDS-polyacrylamide gel (Genscript) and transferred to a nitrocellulose membrane using a wet transfer system (BioRad). Primary antibody mouse anti-tubulin (1:2000) and rabbit anti-FLAG (1:1000) were purchased from Cell Signaling and added overnight at 4°C, followed by three washes in TBS-T (1x TBS-0.1% tween). Secondary antibody IRDye 800CW goat anti-rabbit (1:15,000) and IRDye 680RD goat anti-mouse (1:15,000) were purchased from Li-COR Biosciences and added for 1 h at room temperature, followed by three additional washes in TBS-T. Blots were imaged in IR using a LI-COR Odyssey Blot Imager. Bands were quantified using ImageJ.

**Flow cytometry analysis**

For Panc02 tumor (**Fig. 5**): Panc02 (3 x 10^6^) were subcutaneously injected into the right flank of 6 weeks old female BALB/c mice. Seven days after tumor inoculation, mice were randomized into groups of 3-6 mice and received the following treatment alone or in combination: STF-1623 (0.5, 5, or 50 mg/kg), subcutaneous, on study day 2, day 1-3, or day 1-7; ionizing radiation (15 Gy), day 2. The study was terminated on day 12, and tumors were collected for further flow cytometry analyses. Tumors (<0.8g) were dissected, washed with PBS, and digested using murine tumor dissociation kit (Miltenyi, #130-096-730). Specifically, tumors were minced, added to a Miltenyil C-tube in 3 mL digestion buffer, ran with dissociation program (37_c_m_TDK_1) on GentleMACSTM Octo Dissociator platform. Cells were filtered through a 70 μm stainer, span down at 300 x *g* for 5 minutes, re-suspended in 5 mL FACS buffer (PBS + 2% FBS (Gibco, #10099-141)), and adjusted to 3 x 10^6^ cells per sample. Cells were resuspended in 100 μL FACS buffer with 1 μg/mL Fc-Block (Mouse BD Fc BlockTM, #553141) and incubated at 4 °C for 10 minutes. For surface antibodies Live/dead-efluo780 (eBioscience, #65-0865-14), CD45-BV785 (Biolegend, #103149, clone 30-511), CD3-BUV395 (BD, #740268, clone 17A2), CD4-BUV737 (BD, #612843, clone RM4-5), CD8-PE-eFluor610 (eBiosciences, #61-0081-82, clone 53-6.7), FoxP3-PE (eBiosciences, #12-5773-82, clone FJK-16S), CD335-BV421 (Biolegend, #137612, clone 29A1.4), CD11b-PE-Cy7 (Biolegend, #101216, clone M1/70), F4/80-BV510 (Biolegend, #123135, clone BM8), Gr-1-APC (Biolegend, #108412, clone RB6-8C5), CD206-FITC (Biolegend, #141704, clone C068C2), IA-IE-AF700 (Biolegend, #107622, clone M5/114.15.2), CD69-BV605 (Biolegend, #104530, clone H1.2F3), PD-1-BV650 (BD, #744546, clone J43), cells were stained at 4 °C for 30 minutes in the dark and washed 3 times with FACS buffer. For intracellular markers Ki67 and FoxP3, cells were incubated in 200 μL fixation/permeabilization buffer (eBioscience, #00-5523-00) at room temperature for 30 minutes in the dark followed by 2 mL 1x permeabilization buffer wash twice. Cells were incubated in FoxP3-PE (eBiosciences, #12-5773-82, clone FJK-16S) or Ki67-PerCP-Cy5.5 (Biolegend, # 652424, clone 16A8) antibodies in 100 μL permeabilization buffer at room temperature for 30 minutes in the dark and washed twice with FACS buffer. Samples were analyzed on BD LSRFortessa using Kaluza.

For CT26 tumors (**Fig. S6**), CT26 (1 x 10^5^) were subcutaneously injected into the right flank of 6 weeks old female BALB/c mice. 9 days post tumor inoculation (*pi*), mice were randomized into groups of 10 mice. Animals received STF-1632 (5 or 50 mg/kg) subcutaneously on day 9-11, 16-18 *pi*, anti-PD-1 (10 mg/kg) intraperitoneally on day 9, 12, 16 and 19 *pi*, or a combination of the two. On day 23 *pi*, tumors were collected for further flow cytometry analyses. Following single cell dissociation as described above, cells were first stained with live/dead viability dye along with Fc Block-Tru-Stain Monocyte Blocker, TryStain FcX (anti-mouse CD16/32), each at 1:500 dilution at room temperature for 15 mins. Cells were washed once with FACS wash buffer (Biolegend, Cat# 420201) and then stained with a mixture of antibodies: CD45-PerCP.Cy5.5 (Biolegend, #103132; clone 30-F11), CD3-APC/Fire750 (Biolegend, # 100248, clone 17A2); CD4-BV421 (Biolegend, #100443, clone GK1.5); CD8a-PE.Cy7 (Biolegend, #100722, clone 53-6.7). The staining was carried out at 4°C for 30 minutes followed by 2X washes with FACS wash buffer. The cells were then suspended in 500 µL of FACS wash buffer and an aliquot (200 µL) was transferred to a 96-well round bottom plate to run the samples on the flow cytometer (Attune NxT). Instrument and Compensation settings were achieved using Ultra Compensation Beads (ThermoFisher Scientific, Cat# 01-2222-42). Additional controls such as pooled cell suspensions treated with each antibody as a single stain and Fluorescent Minus One (FMO) controls for each marker were also included. Analysis was done using FlowJo Analysis Software version 10.8.0.

**Immunohistochemistry and histology**

Tumors and lungs underwent formalin-fixed paraffin-embedded (FFPE) preparation and section at 4 μm thickness. To characterize tumor infiltrating lymphocytes, CD8 primary antibody staining (CST, #98941; 1:400) followed by anti-rabbit poly-HRP-IgG (Leica, #DS9800) of tumor samples were performed. Stained sections were scanned with Pannoramic Digital Scanners for 40x magnification. Images were analyzed utilizing the HALO® platform. The whole slide image was analyzed, and areas of large necrosis were excluded. Both total tissue area and the number of IHC-positive cells was counted. IHC scores are presented as CD8^+^ cells/tissue area (mm^2^). To characterize metastases to the lungs, lung PPFE were stained with hematoxylin and eosin (H&E). Metastatic foci were identified by blinded pathologist and quantified (metastasis #/animal, metastasis burden (% area of lung tissue)).

**IFN-γ measurement in tumor and serum using qRT-PCR and ELISA**

EMT6 or MC38 (1 x 10^6^) were subcutaneously injected into female BALB/c or C57BL/6 mice, respectively. When the average tumor volume reached 70-95 mm^3^, one dose of STF-1623 (50 mg/mL) or vehicle (PBS) was injected subcutaneously. Serum and tumor were collected 30 minutes after vehicle injection. Serum was collected 2, 4, 6, 8, 24, 48 hours after STF-1623 injection; tumor was collected 0.25, 0.5, 2, 4, 6, 8, 24, 48 hours after STF-1623 injection. Each group has three mice. Tumor IFN-γ was measured with the ABI ViiA 8 Real Time-PCR system, using Taqman Universal PCR Master Mix (ABI, #4304437) and primer pairs specific for mouse *Ifn-γ* (Mm.PT.58.41152792, IDT, #299208314) and *GapdH* cDNA (Mm.PT.39a1, IDT, #296236491). Average of technical triplicates of each sample were used. Serum IFN-γ was measured using the Meso Scale Discovery® Electrochemiluminescence (MSD-ECL) platform using V-PLEX mouse Cytokine 19-Plex kits (MSD #K15255D-1) containing IFN-γ per the manufacturer’s instructions. For data analysis, the MSD Workbench 4.0 software applies a 4-parameter log fit to the calibrator signal intensities to generate standard curves and determine the lower limit of detection. Sample signal intensities are back interpolated to the standard curves to determine the measured concentrations. Average of technical duplicates of each sample were used.

**Pyrophosphate (PPi) measurement**

Female BALB/c mice with established breast tumor (EMT6 or 4T1) were injected subcutaneously with STF-1632 (50 mg/kg) or vehicle control for seven days (n = 5 mice per group). Mice were sacrificed, and liver, kidney, aorta, and serum were collected. Tissues were homogenized in 50 mM HEPES pH 7.2 to 100 mg/mL. 0.5 μL of serum, kidney, liver, and tumor samples and 1 μL of aorta samples were added to the pyrophosphate reaction following manufacturer instruction (Sigma). The average of two technical duplicates was taken for each sample.

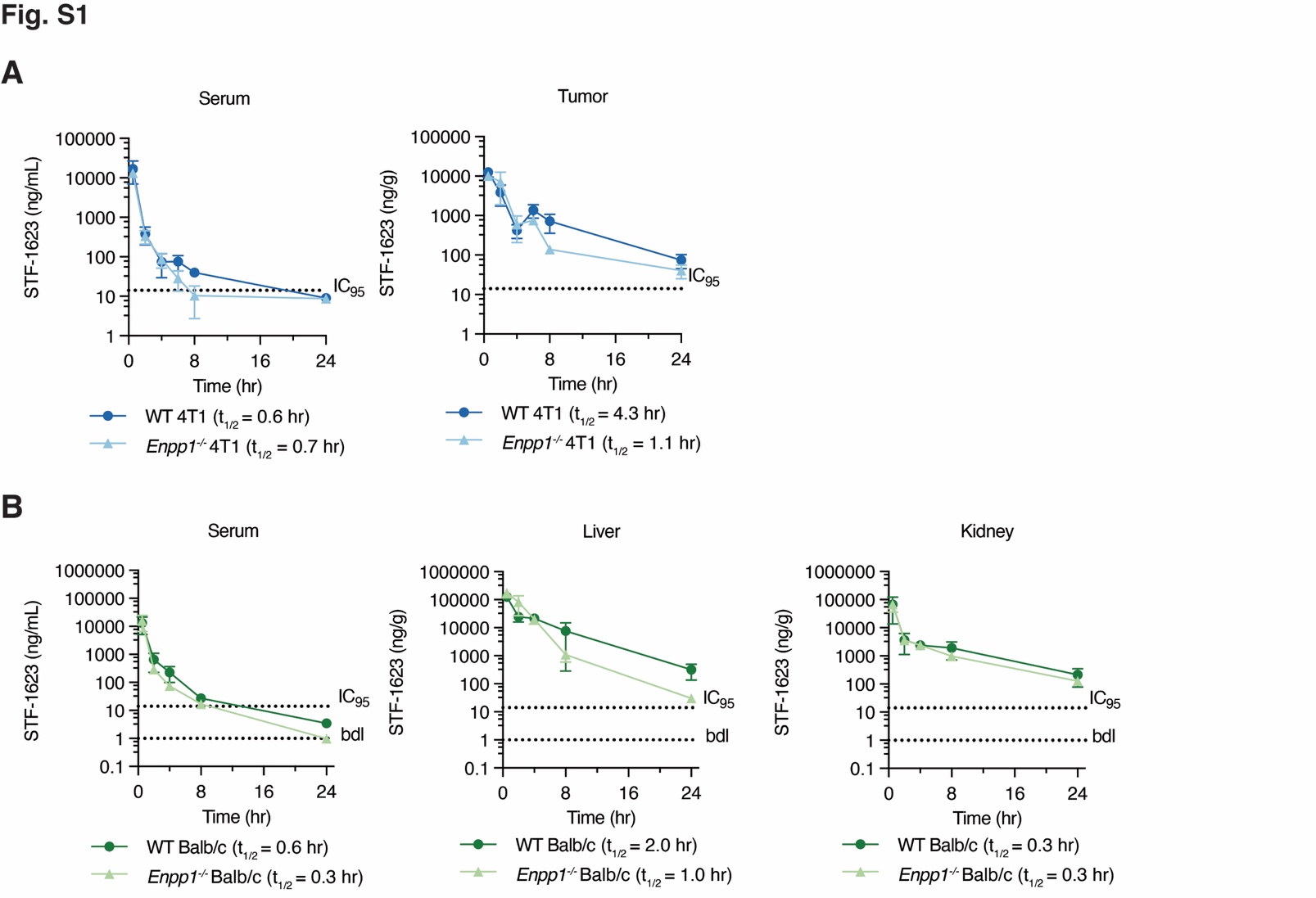

**Fig S1. STF-1623 exhibits target-driven homing to tumors and tissues (related to Figure 1).** (**A**) Concentration of STF-1623 in serum (left) and tumor (right) of WT BALB/cJ mice with established orthotopic WT or *Enpp1^-/-^* 4T1 tumors after one subcutaneous dose of STF-1623 (50 mg/kg). IC_95_ = 14 ng/mL or g. Mean ± SEM is plotted, n = 2 mice for each point except the following, where n = 3: all points at 24 h; n = 1: *Enpp1^-/-^* 4T1 tumor 6h, 8h (point removed as an outlier). (**B**) Concentration of STF-1623 in serum (left), liver (middle), and kidney (right) of WT or *Enpp1^-/-^* BALB/cJ mice after one subcutaneous dose of STF-1623 (50 mg/kg). IC_95_ = 14 ng/mL or g; below detection limit (bdl) = 1 ng/mL or g. Mean ± SEM is plotted, n = 2 mice for each point except the following, where n = 3: *Enpp1^-/-^* mice at 0.5h.

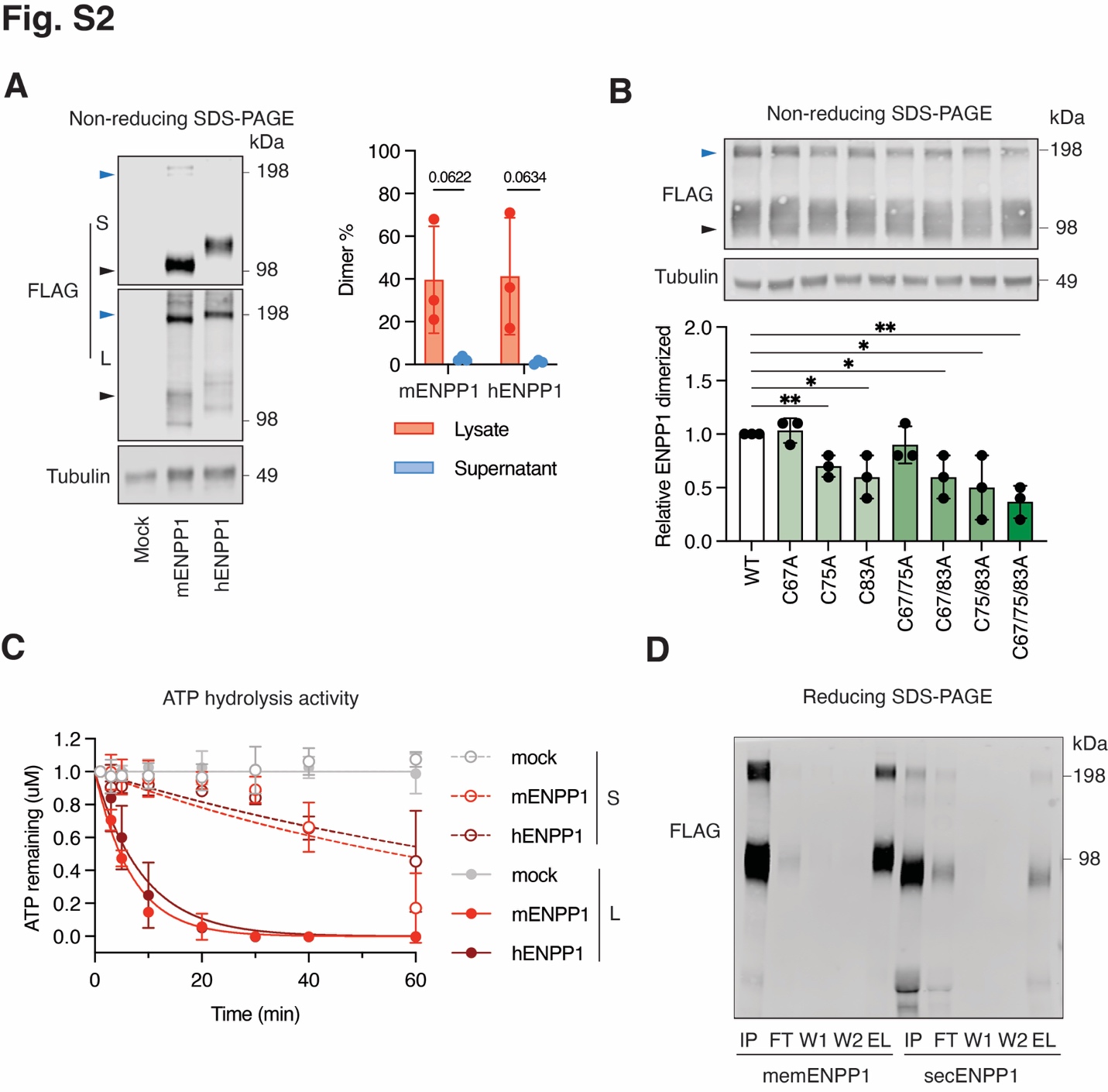

**Fig. S2. Transmembrane and secreted ENPP1 have comparable cGAMP hydrolysis activity (related to Figure 3).** (**A**) Expression of mENPP1, hENPP1 in supernatant and lysate of 293T *ENPP1^-/-^* cells assessed by non-reducing western blotting. Dimer and monomer ENPP1 are indicate with black and blue triangles respectively. Percent dimer is calculated as dimer over the sum of dimer and monomer. Quantification is from three independent experiment, with blots from one representative experiment shown (Full scan of blot available as source data). (**B**) Expression of WT ENPP1 and transmembrane domain cysteine mutants in lysate of 293T *ENPP1^-/-^* cells assessed by reducing non-reducing western blotting. Dimer and monomer ENPP1 are indicate with black and blue triangles respectively. Percent dimer is calculated as dimer over the sum of dimer and monomer. Quantification is from three independent experiment, with blots from one representative experiment shown (Full scan of blot available as source data). (**C**) ATP degradation kinetics of mENPP1 and hENPP1 in lysate and supernatant of 293T *ENPP1^-/-^* cells at pH 9.0. Mean ± SD is plotted, n = 2 technical replicates. Data are fit with one phase decay model with constraints of Y0 at 1 and plateau at 0. (**D**) Expression of purified memENPP1 from lysate and secENPP1 from supernatant of 293T cGAS *ENPP1^-/-^* cells assessed by reducing western blotting. L: lysate; S: supernatant; mENPP1: mouse ENPP1; hENPP1: human ENPP1; IP: input; FT: flow through; W1: wash 1; W2: wash 2; EL: eluent.

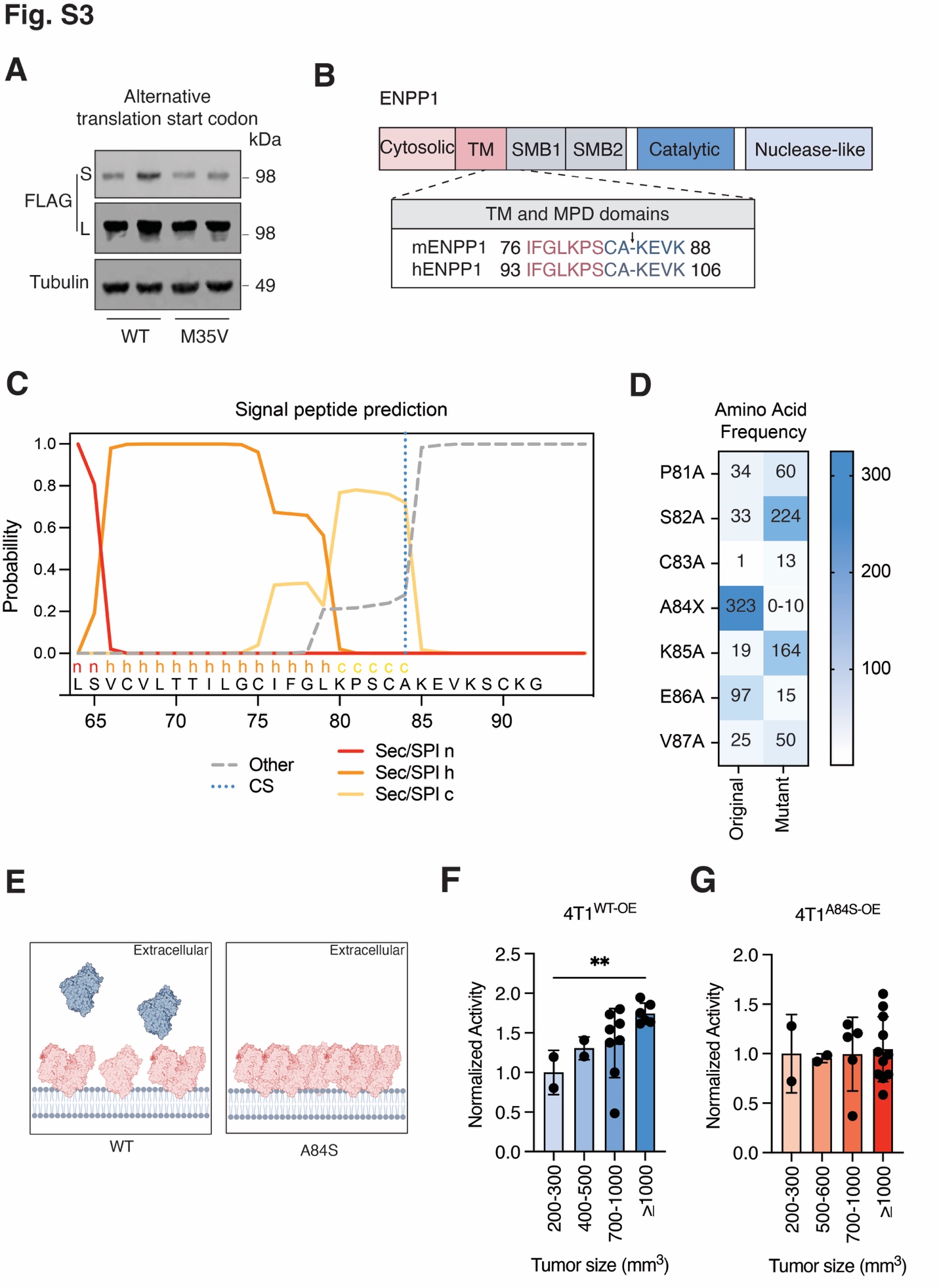

**Fig S3.** **A84S single nucleotide polymorphism increases the membrane and intratumoral retention of ENPP1 (related to Figure 3).** (**A**) Expression of WT and M35V ENPP1 in supernatant and lysate of 293T *ENPP1^-/-^* cells assessed by reducing western blotting. Data are from one experiment (full scan of blot available as source data). (**B**) Protein domains of ENPP1. Zoom in of the region containing the predicted cleavage site (arrow) in mENPP1 (A84/K85) and hENPP1 (A102/K103). (**C**) Prediction of the location of protein cleavage site (CS) and the presence of signal peptides of ENPP1 using SignalP 6.0 (<https://services.healthtech.dtu.dk/services/SignalP-6.0/>). ENPP1 is predicted to be processed through the standard secretory signal peptide transported by the Sec translocon and cleaved by Signal Peptidase I (Sec/SPI) and contains the N-terminal n-region (n), hydrophobic h-region (h), and C-terminal c-region (c) preceding the cleavage site. (**D**) The amino acid frequency of the original and mutated ENPP1 residues at and surrounding the signal peptidase cleavage site of, as predicted by SignalP 6.0. (**E**) Schematic of the impact of membrane retention of the mENPP1 mutant A84S that corresponds to a signal nucleotide polymorphism in human ENPP1 (A102S). (**F**) and (**G**), Relative cGAMP hydrolysis activity calculated by kinetic analysis from sera from mice bearing 4T1^WT-OE^ (F) or 4T1^A84S-OE^ (G) orthotopic tumors of various sizes. 4T1^WT-OE^ or 4T1^A84S-OE^ cells (5 x 10^4^) were orthotopically injected in WT BALB/cJ mice. When tumors reach specific sizes, sera was collected for cGAMP hydrolysis activity analysis. Mean ± SD is plotted. The *P* value was determined by two-sided unpaired *t* test. L: lysate; S: supernatant; TM: transmembrane; MPD: membrane proximal domain; SMB: somatomedin B; mENPP1: mouse ENPP1; hENPP1: human ENPP1. ***P* < 0.01

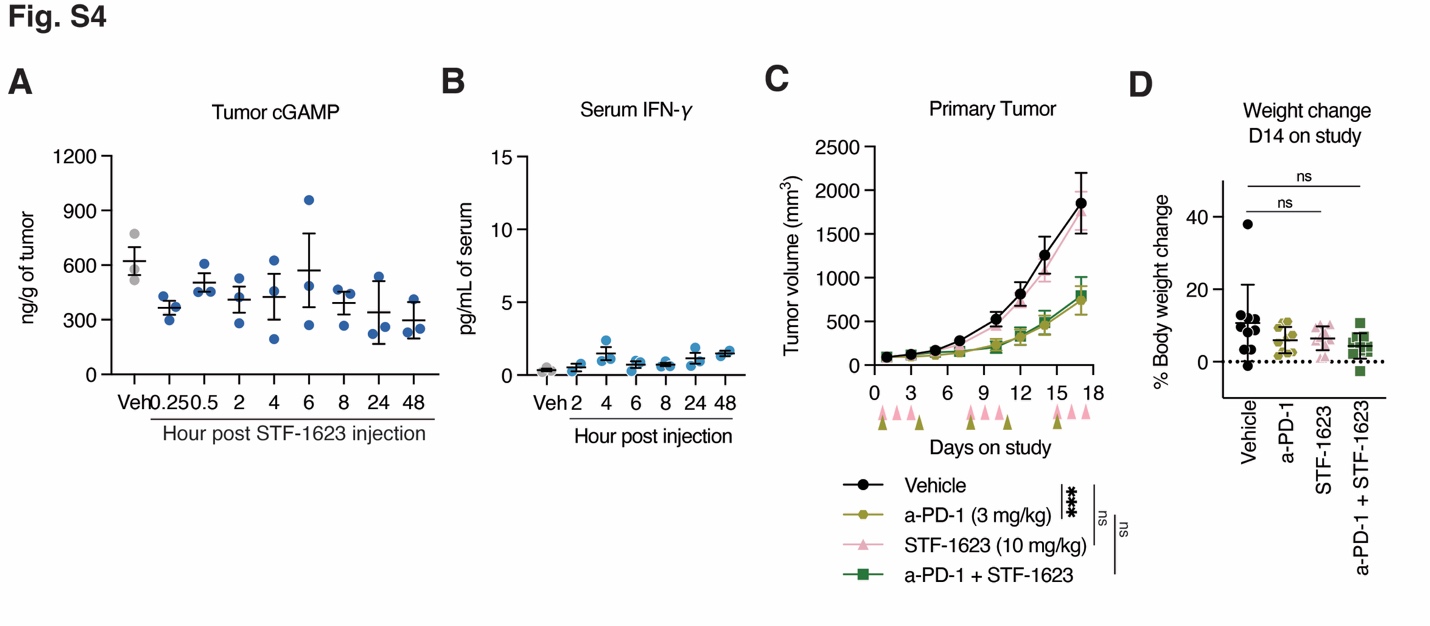

**Fig S4. STF-1623 is not effective towards MC38 colorectal tumor without desired STF-1623 pharmacokinetics and pharmacodynamics profiles (related to Figure 4).** (**A**) cGAMP amount (ng/g tumor) in subcutaneous MC38 tumors at time points indicated after one dose of STF-1623 (50 mg/mL) subcutaneous injection. Mean ± SEM is plotted, n = 3 mice. (**B**) IFN-γ levels (pg/mL serum) in serum of MC38 bearing mice at time points indicated one dose STF-1623 (50 mg/mL) subcutaneous injection. Mean ± SEM is plotted, n = 3 mice (average of technical duplicates). (**C**) MC38 cells (5 x 10^5^) were subcutaneously injected in C57BL/6 mice. When the average tumor size reached ~90 mm^3^, mice were randomized into 10 per group and received anti-PD-1 (a-PD-1), STF-1623 or their combinations at the dosage and frequencies (green and pink triangles respectively) indicated. Mean ± SEM is plotted. The *P* value for tumor volume on day 17 was determined by multiple unpaired *t* test. (**D**), Percent weight change of mice in **c** on day 14 compared to day 1 of the study. Mean ± SD is plotted, n = 10 mice per group. ****P* < 0.001.

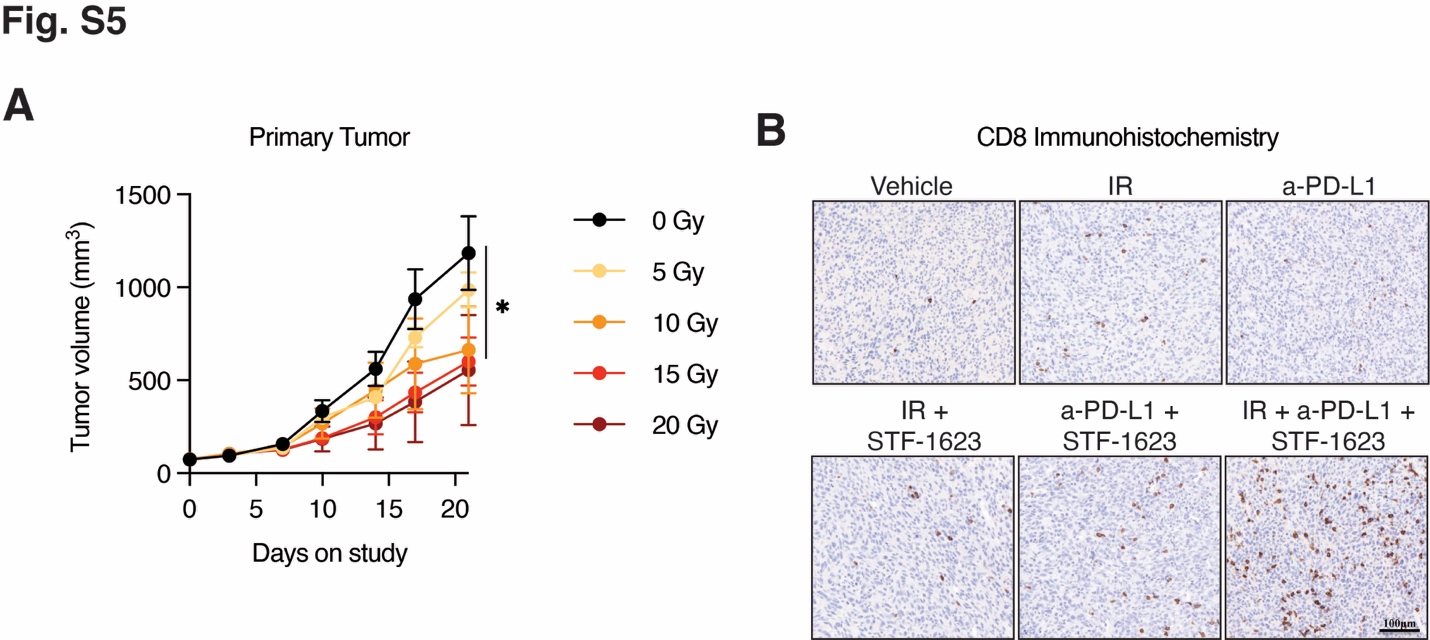

**Fig S5. Characterization of EMT tumors (related to Figure 4).** (**A**) Tumor growth of BALB/c mice bearing established EMT6 subcutaneous tumor receiving one dose of ionizing radiation on study d0 at the dosage indicated. Mean ± SEM is plotted, n = 4 mice per group. (**B**) Representative immunohistochemistry images of CD8+ cells in tumors from mice in **Fig. 4D** (scale bar is 0.1 mm).

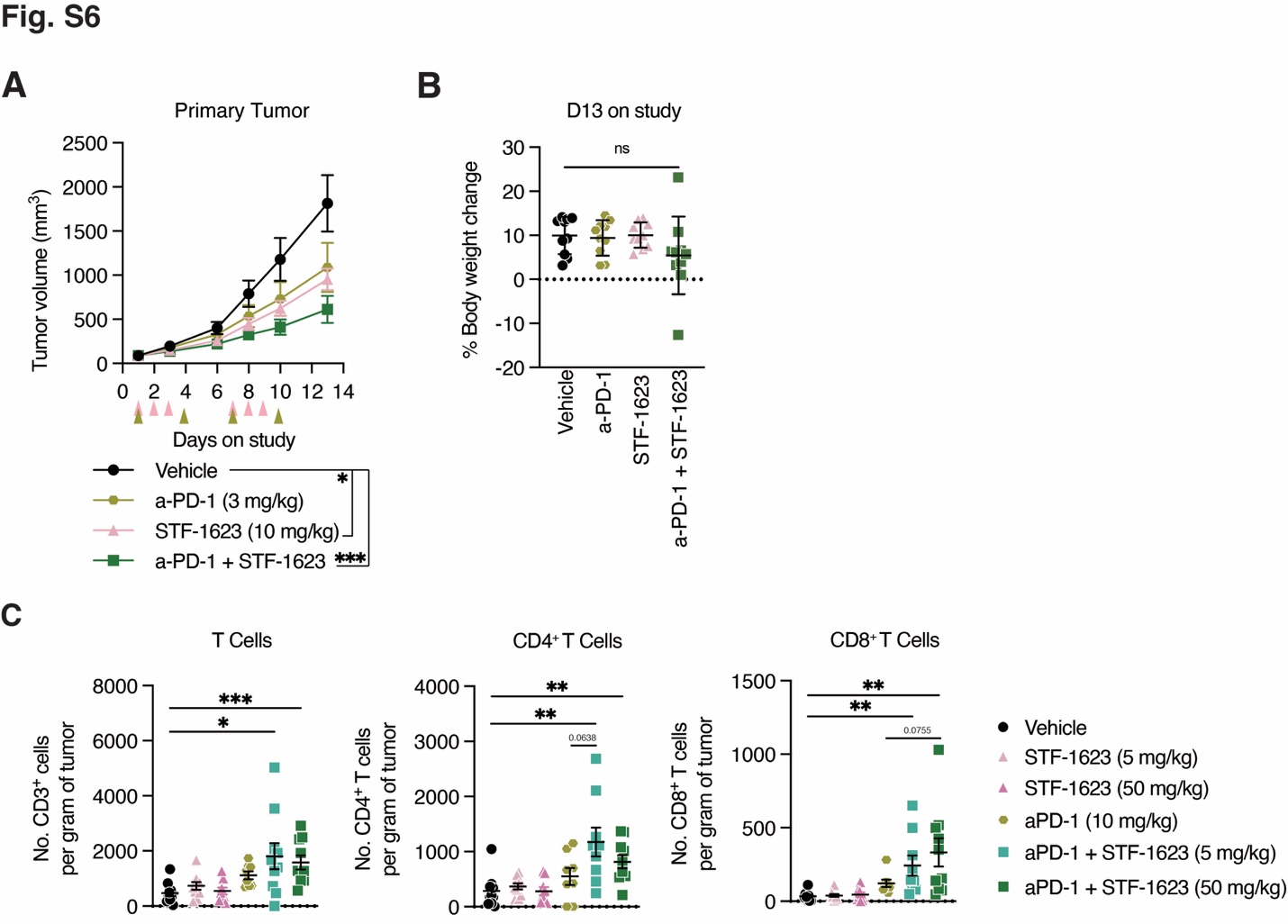

**Fig S6. STF-1623 delays CT26 colorectal tumor growth.** (**A**) CT26 (1 x 10^6^) were subcutaneously injected in BALB/c mice. When the average tumor size reached ~90 mm^3^, mice were randomized and received STF-1632 subcutaneously, anti-PD-1 intraperitoneally, or a combination of the two at dosage and frequency indicated (STF-1623: pink triangle, a-PD-1: green triangle). Mean ± SEM is plotted, n = 10 mice. The *P* value for tumor volume on day 13 was determined by multiple unpaired *t* test. (**B**) Percent weight change of mice in (A) on day 13 compared to day 1 of the study. Mean ± SD is plotted, n = 10 mice. (**C**) CT26 cells (1 x 10^5^) were subcutaneously injected in BALB/c mice. When the average tumor size reached ~100 mm^3^, mice were randomized and received 5 or 50 mg/kg of STF-1632 subcutaneously on day 9-11, 16-18 post inoculation (*pi*), 10 mg/kg anti-PD-1 intraperitoneally on day 9, 12, 16 and 19 *pi*, or a combination of the two. On day 23 of the study, tumors were isolated and processed for flow cytometry. The number of CD3^+^ T cells, CD4^+^CD3^+^ T cells, CD8^+^CD3^+^ T cells per gram of tumors are shown. Mean ± SEM is plotted, n = 10 mice. *P* values were calculated by two-sided unpaired *t* test. **P* < 0.05, ***P* < 0.01, ****P* < 0.001.

**Table S1. Key resource table**

| **REAGENT or RESOURCE** | **SOURCE** | **IDENTIFIER** |
| --- | --- | --- |
| Antibodies | | |
| *InVivo*MAb anti-mouse PD-1 (CD279) | Bio X Cell | Cat# BE0146 |
| *InVivo*MAb anti-mouse PD-L1 (B7-H1) | Bio X Cell | Cat# BE0101 |
| Live/dead-efluo780 | eBioscience | Cat# 65-0865-14 |
| CD45-BV785 (30-511) | BioLegend | Cat# 103149 |
| CD45-PerCP.Cy5.5 (30-F11) | BioLegend | Cat# 103132 |
| CD3-BUV395 (17A2) | BD Biosciences | Cat# 740268 |
| CD3-APC/Fire750 (17A2) | BioLegend | Cat# 100248 |
| CD4-BUV737 (RM4-5) | BD Biosciences | Cat# 612843 |
| CD4-BV421 (GK1.5) | BioLegend | Cat# 100443 |
| CD8-PE-eFluor610 (53-6.7) | eBiosciences | Cat# 61-0081-82 |
| CD8a-PE.Cy7 (53-6.7) | BioLegend | Cat# 100722 |
| FoxP3-PE (FJK-16S) | eBiosciences | Cat# 12-5773-82 |
| CD335-BV421 (29A1.4) | BioLegend | Cat# 137612 |
| CD11b-PE-Cy7 (M1/70) | BioLegend | Cat# 101216 |
| F4/80-BV510 (BM8) | BioLegend | Cat# 123135 |
| Gr-1-APC (RB6-8C5) | BioLegend | Cat# 108412 |
| CD206-FITC (C068C2) | BioLegend | Cat# 141704 |
| IA-IE-AF700 (M5/114.15.2) | BioLegend | Cat# 107622 |
| CD69-BV605 (H1.2F3) | BioLegend | Cat# 104530 |
| PD-1-BV650 (J43) | BD Biosciences | Cat# 744546 |
| Ki67-PerCP-Cy5.5 (16A8) | BioLegend | Cat# 652424 |
| TruStain FcX (anti-CD16/32) | BioLegend | Cat# 101320 |
| CD8α (D4W2Z) XP^®^ Rabbit mAb | Cell Signaling | Cat# 94941 |
| Bond polymer refine detection, anti-rabbit poly-HRP-IgG | Leica | Cat #DS9800 |
| DYKDDDDK (FLAG) Tag Rabbit Ab | Cell Signaling | Cat# 2368 |
| Tubulin (DM1A) Mouse Ab | Cell Signaling | Cat# 3873 |
| IRDye 800CW goat anti-rabbit | LI-COR Biosciences | Cat# 926-32211 |
| IRDye 680RD goat anti-mouse | LI-COR Biosciences | Cat# 926-68070 |
| Chemicals, peptides, and recombinant proteins | | |
| STF-1623 | This paper | N/A |
| 2’3’-cGAMP | (*34*) | N/A |
| [^32^P] 2’3’-cGAMP | (*34*) | N/A |
| Temozolomide | Selleck | Cat# S1237 |
| Mouse ENPP1 (transmembrane and secreted) | This paper | N/A |
| Human ENPP3 | This paper | N/A |
| Human ENPP1 | This paper | N/A |
| Human faux ENPP1 | This paper | N/A |
| anti-DYKDDDDK (FLAG) magnetic agarose | ThermoFischer | Cat# A36797 |
| 3xFLAG peptide | Sigma | Cat# F4799 |
| Critical commercial assays | | |
| Pierce^TM^ 660nm protein assay reagent | ThermoFisher | Cat# 22660 |
| CellTiterGlo | Promega | Cat# G8461 |
| Pyrophosphate assay kit | Sigma-Aldrich | Cat# MAK168 |
| Cytokine 19-Plex kit | MSD | Cat# K15255D-1 |
| Deposited data | | |
| Human faux ENPP1 with STF-1623 structure | This paper | PDB: 9NIR |
| Human ENPP1 (Apo) structure | (*71*) | PDB: 6WFJ |
| Mouse ENPP1 with STF-1084 structure | (*44*) | PDB: 6XKD |
| *Enpp1* and *Enpp3* expression in MC38 | (*45*) | http://tismo.cistrome.org/ |
| SignalP6.0 | (*49*) | https://services.healthtech.dtu.dk/services/SignalP-6.0/ |
| Experimental models: Cell lines | | |
| Human: 293T *ENPP1*^-/-^ cells | (*35*) | N/A |
| Human: Expi293F cells | ThermoFisher | A14527 |
| Mouse: 4T1 | ATCC | Cat# CRL-2539 |
| Mouse: 4T1 *Enpp1^-/-^* cells | (*34*) | N/A |
| Mouse: 4T1 ENPP1^WT-OE^ cells | (*34*) | N/A |
| Mouse: 4T1 ENPP1^T238A-OE^ cells | (*34*) | N/A |
| Mouse: 4T1 ENPP1^A84S-OE^ cells | This paper | N/A |
| Mouse: EMT6 | ATCC | Cat# CRL-2755 |
| Mouse: MC38 | Kerafast | Cat# ENH204 |
| Mouse: Panc02 | ATCC | Cat# CRL-2553 |
| Mouse: CT26 | ATCC | Cat# CRL-2638 |
| Mouse: IVISbrite GL261 Red-FLuc | Revvity | Cat# BW134246 |
| Experimental models: Organisms/strains | | |
| Mouse: C57BL/6 | The Jackson Laboratory | C57BL/6J (000664) |
| Mouse: C57BL/6 | Charles River | C57BL/6NCrl |
| Mouse: BALB/c | The Jackson Laboratory | BALB/cJ (000651) |
| Mouse: BALB/c | Taconic Biosciences | BALB/cAnNTac |
| Mouse: BALB/cJ-*Enpp1^asj-2J/GrsrJ^* | The Jackson Laboratory | JAX: 019107 |
| Oligonucleotides | | |
| See Table S1 for PCR primers | This paper | N/A |
| Recombinant DNA | | |
| pcDNA3-sec-hENPP1-His12 | This paper | N/A |
| pcDNA3-sec-hENPP3-His12 | This paper | N/A |
| pcDNA3-sec-hENPP3-Q244K/E275D-His12 | This paper | N/A |
| pcDNA3-hENPP1-WT-Flag | Genscript | N/A |
| pLenti-CMV-hENPP3-WT-FLAG | (*54*) | N/A |
| pcDNA3-mENPP1-WT-Flag | Genscript | N/A |
| pcDNA3-mENPP1-P81A-Flag | This paper | N/A |
| pcDNA3-mENPP1-S82A-Flag | This paper | N/A |
| pcDNA3-mENPP1-C83A-Flag | This paper | N/A |
| pcDNA3-mENPP1-K85A-Flag | This paper | N/A |
| pcDNA3-mENPP1-E86A-Flag | This paper | N/A |
| pcDNA3-mENPP1-V87A-Flag | This paper | N/A |
| pcDNA3-mENPP1-S82A/K85A-Flag | This paper | N/A |
| pcDNA3-mENPP1-A84S-Flag | This paper | N/A |
| pcDNA3-mENPP1-A84G-Flag | This paper | N/A |
| pcDNA3-mENPP1-A84V-Flag | This paper | N/A |
| pcDNA3-mENPP1-A84T-Flag | This paper | N/A |
| pcDNA3-mENPP1-A84P-Flag | This paper | N/A |
| pcDNA3-mENPP1-A84K-Flag | This paper | N/A |
| pcDNA3-mENPP1-A84D-Flag | This paper | N/A |
| pcDNA3-mENPP1-A84F-Flag | This paper | N/A |
| pcDNA3-mENPP1-C67A-Flag | This paper | N/A |
| pcDNA3-mENPP1-C75A-Flag | This paper | N/A |
| pcDNA3-mENPP1-C67/75A-Flag | This paper | N/A |
| pcDNA3-mENPP1-C67/83A-Flag | This paper | N/A |
| pcDNA3-mENPP1-C75/83A-Flag | This paper | N/A |
| pcDNA3-mENPP1-C67/75/83A-Flag | This paper | N/A |
| pcDNA3-mENPP1-M35V-Flag | This paper | N/A |
| pcDNA3-hENPP1-Flag | Genscript | N/A |
| pLenti-CMV-GFP-Puro | Addgene | RRID: Addgene_17448 |
| pLenti-CMV-mENPP1-A84S-GFP-Puro | This paper | N/A |
| Software and algorithms | | |
| Prism 9.1.0 | Graphpad | https://www.graphpad.com/scientific-software/prism/ |
| ImageJ 2.0.0 | Schneider | https://imagej.nih.gov/ij/ |
| Pymol |  | https://www.pymol.org/2/ |
| FlowJo 10.8.0 | FlowJo, LLC | https://www.flowjo.com/ |
| StudyDirectorTM 3.1.399.19 |  |  |

**Table S2. Oligonucleotide Sequences**

| **Name** | **Sequence (5’->3’)** |
| --- | --- |
| Q244K_hENPP3_fwd | CACTTTCTTCAAAGGAAAAAAATAATCCAGCCTGGTGGCATG |
| Q244K_hENPP3_rev | CATGCCACCAGGCTGGATTTTTTTGTTCCTTTGAAGAAAGTG |
| E275D_hENPP3_fwd | CTTTTGGCCCGGATCAGACGTGGCTATAAATGGCTCCTTTC |
| E275D_hENPP3_rev | GAAAGGAGCCATTTATAGCCACGTCTGATCCGGGCCAAAAG |
| pLenti_mENPP1_fwd | gccatccacgctgttttgacctccatagaagacaccgactctagaGATCCGCCACCATGGAGC |
| pLenti_mENPP1_rev | aacagctcctcgcccttgctcaccatggtggcgaccggtggatccCAGAATTCGTCTTCTTGGCTGAAGATTG |
| P81A_mENPP1_fwd | TGTATATTTGGGTTGAAAgCAAGCTGCGCCAAAGAAGT |
| P81A_mENPP1_rev | ACTTCTTTGGCGCAGCTTGCTTTCAACCCAAATATACA |
| S82A_mENPP1_fwd | TATTTGGGTTGAAACCAgcCTGCGCCAAAGAAGT |
| S82A_mENPP1_rev | ACTTCTTTGGCGCAGGCTGGTTTCAACCCAAATA |
| C83A_mENPP1_fwd | GGGTTGAAACCAAGCgcCGCCAAAGAAGTAAA |
| C83A_mENPP1_rev | TTTACTTCTTTGGCGGCGCTTGGTTTCAACCC |
| K85A_mENPP1_fwd | ggttgaaaccaagctgcgccgcagaagtaaaaagttgcaaag |
| K85A_mENPP1_rev | ctttgcaactttttacttctgcggcgcagcttggtttcaacc |
| E86A_mENPP1_fwd | AAACCAAGCTGCGCCAAAGcAGTAAAAAGTTGCAAAGG |
| E86A_mENPP1_rev | CCTTTGCAACTTTTTACTGCTTTGGCGCAGCTTGGTTT |
| V87A_mENPP1_fwd | CAAGCTGCGCCAAAGAAGcAAAAAGTTGCAAAGGCC |
| V87A_mENPP1_rev | GGCCTTTGCAACTTTTTGCTTCTTTGGCGCAGCTTG |
| S82A/K85A_mENPP1_fwd | TATTTGGGTTGAAACCAGCCTGCGCCGCAGAAGT |
| S82A/K85A_mENPP1_rev | ACTTCTGCGGCGCAGGCTGGTTTCAACCCAAATA |
| A84S_mENPP1_fwd | tgggttgaaaccaagctgcTCCaaagaagtaaaaagttgc |
| A84S_mENPP1_rev | gcaactttttacttctttGGAgcagcttggtttcaaccca |
| A84G_mENPP1_fwd | tgggttgaaaccaagctgcGGCaaagaagtaaaaagttgc |
| A84G_mENPP1_rev | gcaactttttacttctttGCCgcagcttggtttcaaccca |
| A84V_mENPP1_fwd | tgggttgaaaccaagctgcGTCaaagaagtaaaaagttgc |
| A84V_mENPP1_rev | gcaactttttacttctttGACgcagcttggtttcaaccca |
| A84T_mENPP1_fwd | tgggttgaaaccaagctgcACCaaagaagtaaaaagttgc |
| A84T_mENPP1_rev | gcaactttttacttctttGGTgcagcttggtttcaaccca |
| A84P_mENPP1_fwd | tgggttgaaaccaagctgcCCCaaagaagtaaaaagttgc |
| A84P_mENPP1_rev | gcaactttttacttctttGGGgcagcttggtttcaaccca |
| A84K_mENPP1_fwd | tgggttgaaaccaagctgcAAAaaagaagtaaaaagttgc |
| A84K_mENPP1_rev | gcaactttttacttctttTTTgcagcttggtttcaaccca |
| A84D_mENPP1_fwd | tgggttgaaaccaagctgcGACaaagaagtaaaaagttgc |
| A84D_mENPP1_rev | gcaactttttacttctttGTCgcagcttggtttcaaccca |
| A84F_mENPP1_fwd | tgggttgaaaccaagctgcTTCaaagaagtaaaaagttgc |
| A84F_mENPP1_rev | gcaactttttacttctttGAAgcagcttggtttcaaccca |
| C67A_mENPP1_fwd | CGCTGGTTTTGTCAGTAgcTGTGCTAACAACAATTC |
| C67A_mENPP1_rev | GAATTGTTGTTAGCACAGCTACTGACAAAACCAGCG |
| C75A_mENPP1_fwd | CTAACAACAATTCTTGGTgcTATATTTGGGTTGAAACCAAGC |
| C75A_mENPP1_rev | GCTTGGTTTCAACCCAAATATAGCACCAAGAATTGTTGTTAG |
| M35V_mENPP1_fwd | ctgctcgcgcccgtggacctaggag |
| M35V_mENPP1_rev | ctcctaggtccacgggcgcgagcag |

**Table S3. Data collection and refinement statistics**

|  | Human ENPP3 (Q244K, E275D)-Inhibitor complex |
| --- | --- |
| **Data collection** |  |
| X-Ray Source  Wavelength (Å)  Space group | SSRL BL12-2  0.76910  P4_3_2_1_2 |
| Cell dimensions |  |
| *a*, *b*, *c* (Å) | 72.93, 72.93, 382.73 |
| α, β, γ () | 90.00, 90.00, 90.00 |
| Matthews coefficient (Å^3^/Da)^a^  Solvent content (%)  Wilson B value (Å^2^)  Mosaicity/Anisotropy  Resolution (Å)^b^ | 2.73  54.9  42.3  0.28/0.72  37.45(2.70) |
| *R*_merge_^c^ | 0.209(0.852) |
| *I* / σ*I* ratio^d^ | 5.4(1.8) |
| Completeness (%)^e^ | 96.7(98.4) |
| Reflections (total/unique)  Redundancy^f^ | 121,085(28,447)  4.3(4.2) |
| **Refinement** |  |
| Resolution (Å) | 30.68-2.70 |
| No. reflections/test set | 26,944/1,475 |
| *R*_work_ / *R*_free_^g^ | 27.0/33.3 |
| Mean B value (Å^2^)  F_obs_-F_calc_ correlation^h^  No. atoms | 49.0  0.87 |
| Protein | 6,567 |
| Ligand/ion | 24 (inhibitor); 5 (ions: 2x Zn^2+^, 2x Cl^-^, 1x Ca^2+^); 315 (glycan) |
| Water | 52 |
| *B*-factors |  |
| Protein | 47.5 |
| Ligand/ion | 44.7 (inhibitor); 37.9 (ions); 92.6 (glycan) |
| Water | 24.1 |
| Deviation from ideality (Rmsd values)  Ramachandran statistics^i^ | 0.002 Å (bond length), 1.062° (bond angle) |
| Most favored/allowed regions (%) | 99.1 (802 over 810) |
| Disallowed regions (%) | 0.9 (8 over 810) |
| PDB code | 9NIR |

^a^Ratio of the volume of the asymmetric unit to the molecular weight of all protein in the asymmetric unit

^b^Value in parentheses is for the highest-resolution shell: 2.70 - 2.77 Å.

^c^Reliability factor for symmetry-related reflections calculated as: *R*_merge_ = Σ_hkl_ Σj=1 to N | I_hkl_ – I_hkl_ (j) | / Σ_hkl_ Σj=1 to N I_hkl_ (j), where N is the redundancy of the data. In parentheses, the cumulative value at the highest-resolution shell

^d^Ratio of mean intensity to the mean standard deviation of the intensity over the entire resolution range

^e^Fraction of measured reflections to possible observations at the resolution range

^f^Number of measurements of individual, symmetry unique reflections

^g^Average deviation between the observed and calculated structure factors calculated as: *R*_work_ = Σ_hkl_ ||F_obs_| - |F_calc_|| / Σ_hkl_ |F_obs_|, where the F_obs_ and F_calc_ are the observed and calculated structure factor amplitudes of reflection hkl. *R*_free_ is equal to *R*_factor_ but for a randomly selected 5.0 % subset of the total reflections that were held aside throughout refinement for cross-validation

^h^Correlation coefficient between observed and calculated structure factor amplitudes

^i^According to Rampage implemented in CCP4

**Table S4. HPLC-MS parameter details**

| **Liquid chromatography (Shimadzu UFLC XR) conditions** | | |
| --- | --- | --- |
| Compound | STF-1623 | I.S. (Carbamazepine) |
| Column | Phenomenex Gemini 3µ C6-phenyl, 50x2.0mm | |
| Mobile phase: | A: Water with 0.1% Formic Acid  B: Acetonitrile with 0.1% Formic Acid | |
| Flow rate (ml/min) | 0.35 | |
| Temperature (°C) | 35 | |
| Injection volume (µl) | 10 | |

| **Gradient elution conditions** | | |
| --- | --- | --- |
| Time (min) | Mobile phase A (%) | Mobile phase B (%) |
| 0.2 | 90 | 10 |
| 0.5 | 90 | 10 |
| 2.0 | 5 | 95 |
| 3.0 | 5 | 95 |
| 4.0 | 90 | 10 |
| 5.9 | 90 | 10 |

| **Mass spectrometry (API5500) conditions** | | |
| --- | --- | --- |
| Compound | STF-1623 | I.S. (Carbamazepine) |
| MRM(+) | 352.1/334.2 | 237.2/194.1 |
| Collision Gas | 7 | |
| Curtain GAS | 35 | |
| Ion Source Gas1 | 55 | |
| Ion Source Gas2 | 50 | |
| Ion Spray Voltage | 5500 | |
| Temperature (°C) | 550 | |
| Collision Energy | 36 | 26 |
| Declustering Potential | 34 | 136 |
| Entrance Potential | 10 | |
| Collision Cell Exit Potential | 14 | |

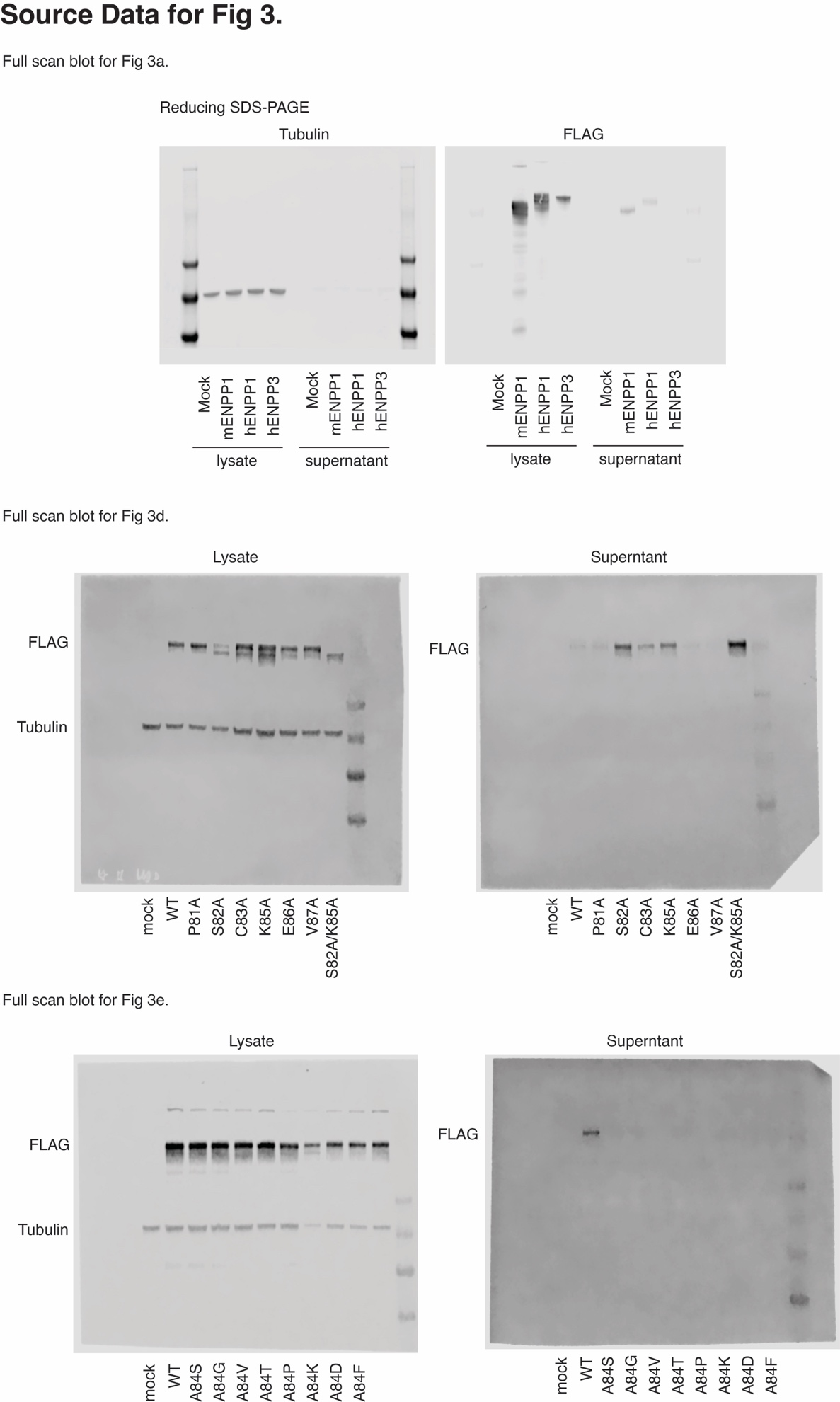

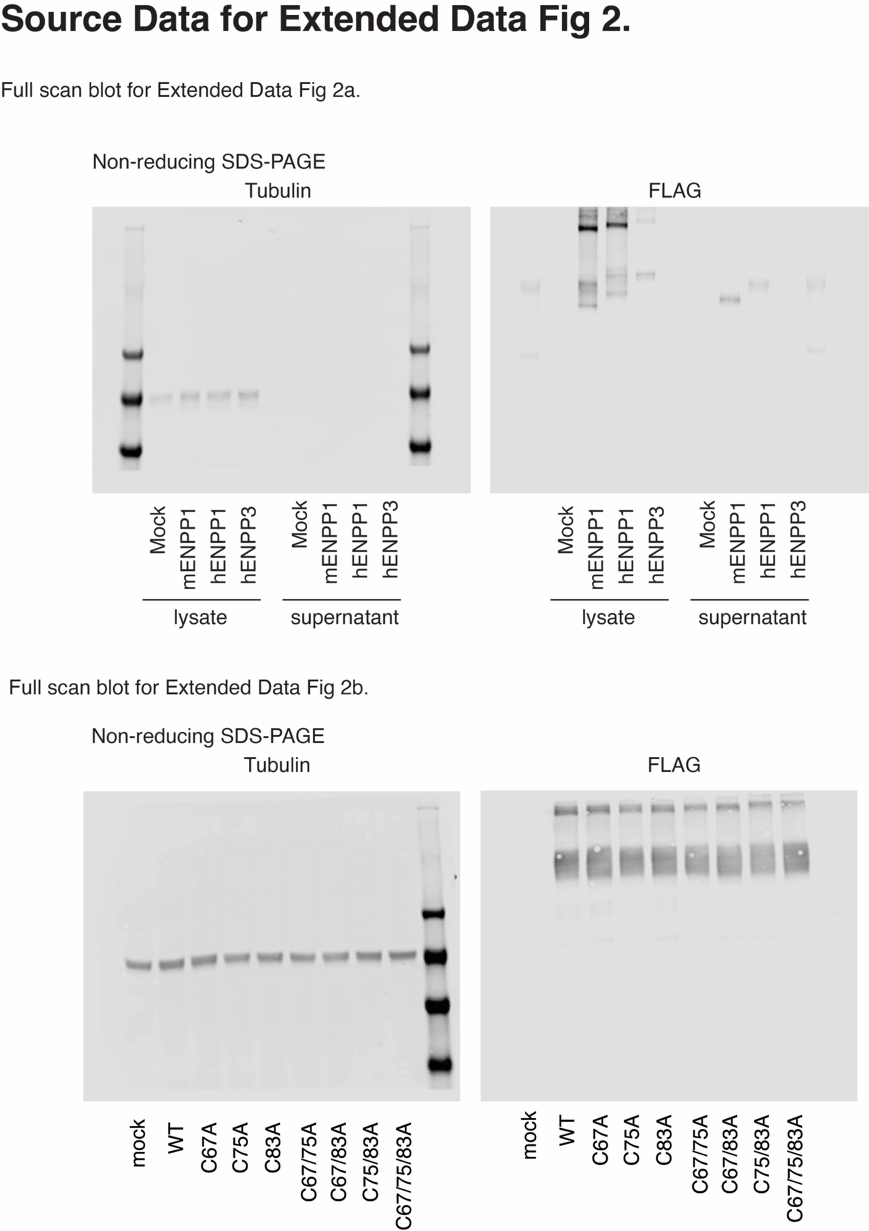

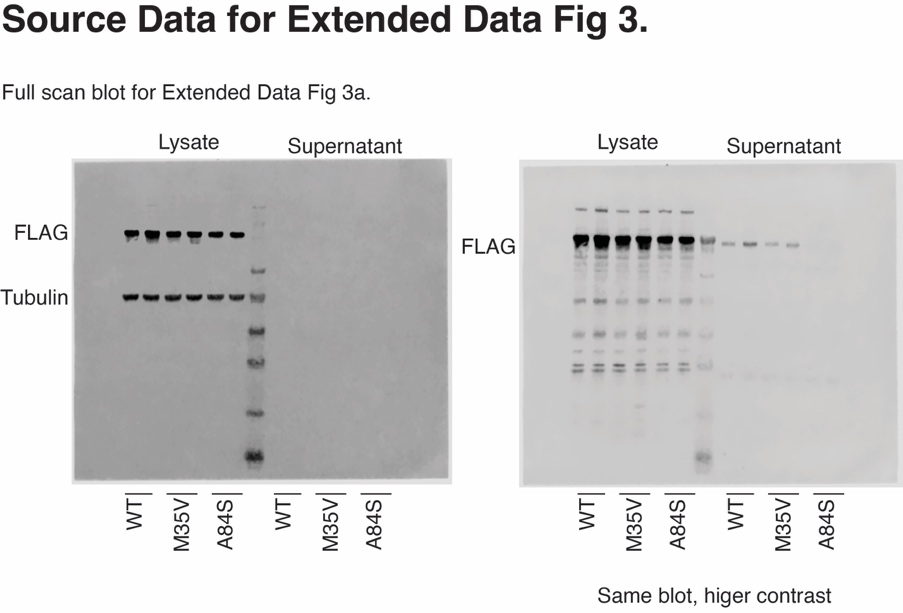
